# Structure of the Disulfide-rich Modules of a Striking Tandem Repeat Protein, Avian Cysteine-Rich Eggshell Membrane Protein

**DOI:** 10.1101/2025.09.16.676614

**Authors:** Somayeh Zeinalilathori, Ryan W. Russell, Jean L. Ross, Shannon Modla, Jeffrey L. Caplan, Tatyana Polenova, Colin Thorpe

## Abstract

Avian eggshell membrane (ESM) is fabricated within the isthmus region of the oviduct and is comprised of three juxtaposed, predominantly proteinaceous layers lying between egg white and the calcified shell. The limiting membrane is less than 0.5 µm in thickness and forms the osmotic barrier for the egg. This first layer provides the foundation for the successive deposition of two mats of protein fibers. Fibers from both inner and outer layers appear to have similar amino acid compositions and are notably disulfide-rich (comprising about 10% Cys). ESM has been utilized in a wide variety of applications, including nutraceutical supplements, tissue engineering, and nanofabrication, and yet fundamental questions concerning protein composition, fiber structure, and membrane assembly remain to be resolved. We previously identified an abundant disulfide-rich structural protein in chicken ESM fibers (Cysteine-rich eggshell membrane protein; CREMP) that contains multiple tandemly repeated modules. In this work, we determine a structural model for four consecutive 2-disulfide containing CREMP modules using a variety of two- and three-dimensional solution NMR experiments. CREMP modules feature an N-terminal loop region positioned above a small beta hairpin that is stabilized by a conserved pattern of disulfide bridges between Cys1-3 and Cys2-4. While the individual CREMP modules are highly ordered, the lack of long-range inter-module restraints suggests an extended structure connected by flexible linkers. Finally, the structural information obtained in this work is considered in the context of full-length CREMP proteins and compared to two other structural proteins that contain multiple tandem repeats of 2-disulfide modules.

**Statement of Importance and Impact:** Under the shell of an avian egg is a tough, paper-thin, three-layered protein-based structure called the eggshell membrane. This membrane has a range of medical, biomaterial and analytical applications, and yet fundamental questions concerning protein composition and assembly remain unresolved. Here, we address the 3-dimensional structure of part of a major membrane component, cysteine-rich eggshell membrane protein. The work provides new insight into the eggshell membrane and suggests an additional avenue for de-novo protein design.

## INTRODUCTION

The eggshell membrane (ESM) of avian eggs is a tripartite proteinaceous structure that forms the foundation for the deposition of the calcified shell (Burley & Vadehra, 1989). Although this membrane has a wide range of current and anticipated applications (including biomedical, nutraceutical, dye, and heavy metal sequestration, and as a template for tissue engineering and nanofabrication (Balaz et al., 2021; Kulshreshtha et al., 2022; Mensah et al., 2022; Mittal et al., 2016; Shi et al., 2021), it remains remarkably poorly understood at a molecular level.

Figure 1 shows an image of chicken ESM using serial block-face scanning electron microscopy (SBF-SEM; see Methods). A supplementary video (video S1) provides a striking additional 3D perspective. The structure is dominated by proteinaceous fibers that are each constructed from a central core with an overcoat of mantle material that is less electron dense when samples are stained using typical electron-contrast reagents (Bellairs & Boyde, 1969; Simons & Wiertz, 1963; Tan et al., 1992). Fabrication of ESM starts with the creation of a thin proteinaceous film (of <0.5 µm) over egg white that represents the osmotic barrier and provides the final physical restraint from microbial attack (Bellairs & Boyde, 1969; Hincke et al., 2000; Tan et al., 1992). This limiting membrane is fused to the so-called ESM inner membrane (Figure 1 and video S1) by what appears to be an extension of the mantle material (Bellairs & Boyde, 1969). This inner ESM membrane is ∼20 µm thick and is formed from relatively thin fibers with widths of ∼1-3 µm (Bellairs & Boyde, 1969; Tan et al., 1992). The adjoining ∼60 µm thick outer ESM is fabricated from noticeably wider fibers (particularly evident in the expanded ESM section shown in Figure S1 and video S2). Outer and inner membranes are readily separated by exploiting their segregation at the air sac of a chicken egg and have comparable amino acid compositions (Burley & Vadehra, 1989). The images and supplementary videos presented here suggest that ESM forms a comprehensively interconnected architecture secured by a contiguous overcoat of mantle material (Bellairs & Boyde, 1969; Hoffer, 1971; Wangensteen & Weibel, 1982). The net result is a robust, flexible layered assembly that presents, at one face, the critical osmotic barrier retaining the egg white, and on the other, the site of mineralization within the shell gland of the avian oviduct.

**Figure 1.**
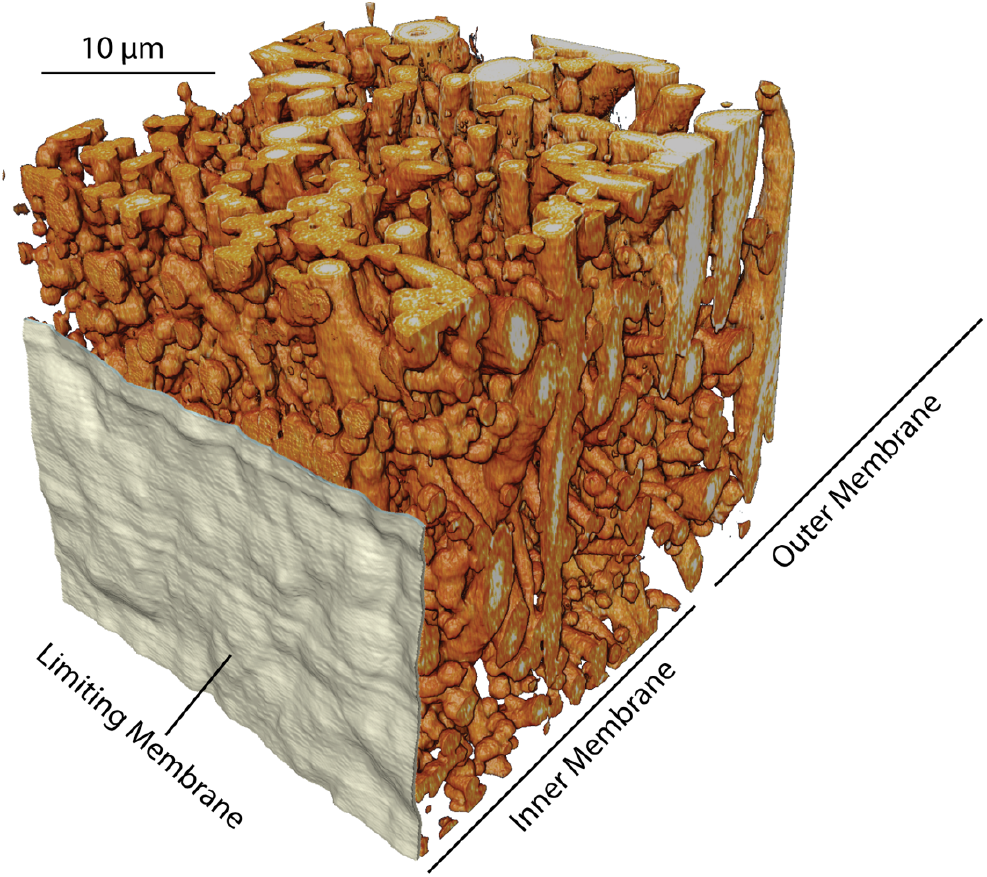
Serial block-face electron microscope image of eggshell membrane. ESM was fixed and then stained with osmium tetroxide, uranyl acetate, and lead aspartate (see Methods (Hua et al., 2015)). 3D reconstructions of 50 nm slices utilized Amira 3D software. The limiting membrane is shown in white and is omitted in the upper portion of the image to show the underlying contact surface at the interfacial region of the inner eggshell membrane. A more extensive static view of ESM is shown in Figure S1. Accompanying videos (video S1 and video S2) provide more comprehensive 3D representations of ESM architecture.

Although the morphology of the ESM has been addressed in classic monographs and papers stretching back decades (Romanoff & Romanoff, 1949), the precise composition of this largely proteinaceous material (∼93% of ESM dry weight (Burley & Vadehra, 1989)) remains cryptic. While the presence of multiple disulfides would not represent a significant impediment to separating the ESM into its constituent proteins, the additional abundance of lysine post-translational modifications is a major challenge. Lysine crosslinks in ESM range from simple Schiff base adducts between the aldehyde side chain of a lysine and the unmodified amino group of a neighboring lysine residue, to the largely intractable desmosine and isodesmosine crosslinks formed from the oxidative condensation of 4 lysine side chains (Burley & Vadehra, 1989; Crombie et al., 1981; Leach et al., 1981). This extensive derivatization of Lys residues, the presence of additional post-translational modifications including hydroxylation and glycosylation reactions (Candlish & Scougall, 1969; Wong et al., 1984), and the likely entrapment/association of incidental non-structural globular proteins within the ESM fibrous network, complicate efforts to apply quantitative proteomics to identify the major structural proteins forming this complex biomaterial.

The early recognition of desmosine and isodesmosine in amino acid hydrolysates of the ESM (Leach et al., 1981; Starcher & King, 1980) and the finding that collagen immunoreactive material is present in the fibers (Arias et al., 1991; Wong et al., 1984) has led to the still-pervasive claim that ESM is “largely collagen”. This assertion is highly improbable. Several independent amino acid analyses of ESM (summarized in (Kodali et al., 2011)) give an average of 10.1 ± 0.7 % Cys, whereas avian collagens are notable for their paucity of this amino acid (chicken collagens I, V, and X found in ESM (Du et al., 2015; Wong et al., 1984) average about 0.8% Cys). Even if it were assumed that just half of ESM proteins were collagen, the remainder would require an aggregate Cys content approaching 20%. No evidence for the existence of such exceptionally Cys-rich proteins has emerged from proteomic studies of the ESM (Ahmed et al., 2017; Ahmed et al., 2019; Kodali et al., 2011; Ma et al., 2025).

In 2011, our laboratory reported the presence of peptides from partial chicken ESM digests that were similar to Spore Coat Protein from the slime molds *Dictyostelium discoideum* and *Polysphondilium pallidum* (West, 2003). Kodali et al. (Kodali et al., 2011) named these proteins Cysteine-Rich Eggshell Membrane Proteins (CREMP). Chicken CREMP proteins have Cys contents of 12-13% and feature runs of consecutive 2-disulfide motifs that extend over more than two thousand residues (Figure 2A and supplemental Figure S2A and S2B) (Kodali et al., 2011). Towards the C-terminus, there are three von Willebrand Factor type D domains (VWFD) and two adjacent zonadhesin-like cysteine-rich C8 domains. The notable sequence similarity of these CREMP modules is illustrated in Figure 2B using 20 consecutive modules (45-64) out of a total of 81 CREMP motifs in NCBI RefSeq XP_040551000 (annotated as Zonadhesin isoform X2). A WebLogo (Crooks et al., 2004) representation of these 20 consecutive modules (Figure 2C) and all CREMP modules (Figure S2C) illustrates the striking conservation of cysteine spacing within these tandem repeat CREMP proteins (Kodali et al., 2011). We note that the 2-disulfide modules that form the focus of this study have sometimes been annotated as Follistatin-like N-terminal domain. For consistency with our earlier work, we will continue to describe them as CREMP modules.

**Figure 2.**
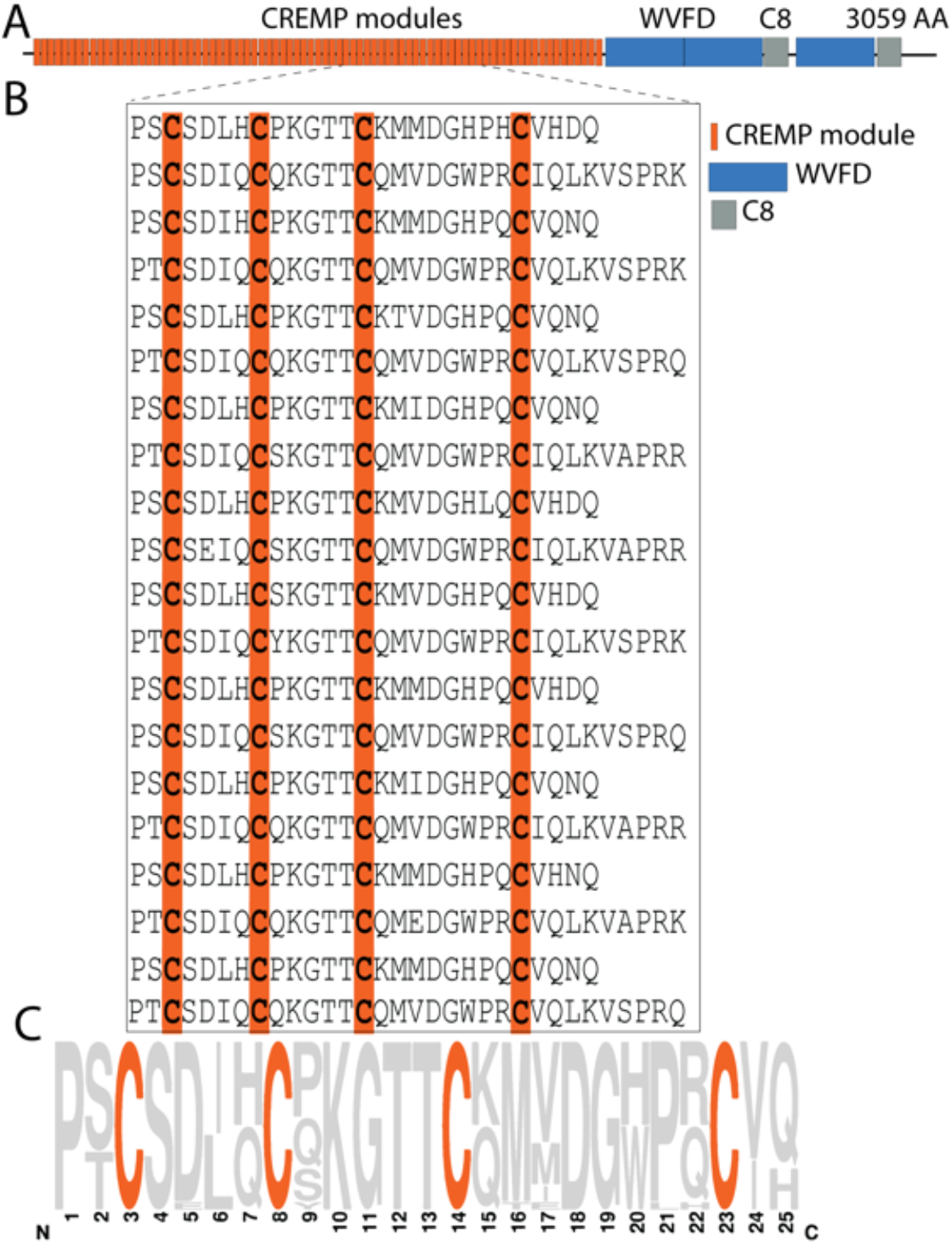
Schematic domain organization and CREMP modules. **Panel A**. The orange bar shows that 2254 of the 3059 amino acid residues in RefSeq XP_040551000.1 represent 81 consecutive CREMP modules (see the Text). Three von Willebrand Factor type D domains (VWFD: dark blue) and the two zonadhesin-like cysteine-rich C8 domains (grey) are located towards the C-terminus (Lu et al., 2020b; Sigrist et al., 2013). **Panel B** depicts 20 consecutive CREMP modules starting from PRO1217. Here, we arbitrarily represent a module starting two residues before the first Cys residue. In the full-length CREMP sequence (Figure S2B), the 4^th^ Cys of one module is separated from the 1^st^ Cys of the next module by between 5 and 17 residues. In Panel B (modules 45-64), this spacing alternates between 6 and 11 residues, and this feature is found in the structure of the 4 consecutive modules examined in this work. A Weblogo (Crooks et al., 2004) depiction of these 20 CREMP modules trimmed to 25 residues long, starting 2 residues before Cys1 of each module, is shown in Panel C. In the full-length sequence (Figure S2B), 78 of 81 CREMP modules show conserved Cys placements at positions 3, 8, 14, and 23. Each of the remaining 3 modules has one amino acid removed from the intra-cysteine spacer regions.

CREMP proteins are likely to be a major component of ESM (Kodali et al., 2011). Thus, comparing the amino acid composition between candidate proteins and ESM hydrolysates using Euclidean distance reveals that chicken CREMP provides a better match to the ESM than collagen or any other single candidate protein, especially considering their cysteine content (Kodali et al., 2011). The likely abundance of CREMP proteins in ESM suggests that they play important roles in the formation of this resilient, multilayered assembly. As a first step towards a structural and contextual understanding of these remarkable CREMP proteins, we report the solution NMR structural model of 4 consecutive CREMP modules. Finally, for perspective, we briefly consider the CREMP model structure in the context of other disulfide-rich tandem repeat structural proteins.

## RESULTS AND DISCUSSION

### Structural characterization of 4-module (4-mod) CREMP

Repeated crystallization trials (either in-house or at the HWI National High-throughput Crystallization Center (Lynch et al., 2023)), using either the TEV-cleaved 4-mod CREMP or the MBP-fused construct, were unsuccessful. Both samples appear highly prone to phase separation/coacervation without subsequent crystal formation. In contrast, 4-mod CREMP proved robust in solution and amenable to solution NMR for structural characterization. Figure 3A shows a 2D ^1^H-^15^N-HSQC spectrum of 4-mod CREMP. The spectrum is surprisingly well-dispersed, given the high level of sequence degeneracy. 108 resolved peaks out of 109 non-proline residues were seen, indicating that 4-mod CREMP is well-ordered and monomeric in solution, consistent with our previous study (Kodali et al., 2011).

**Figure 3.**
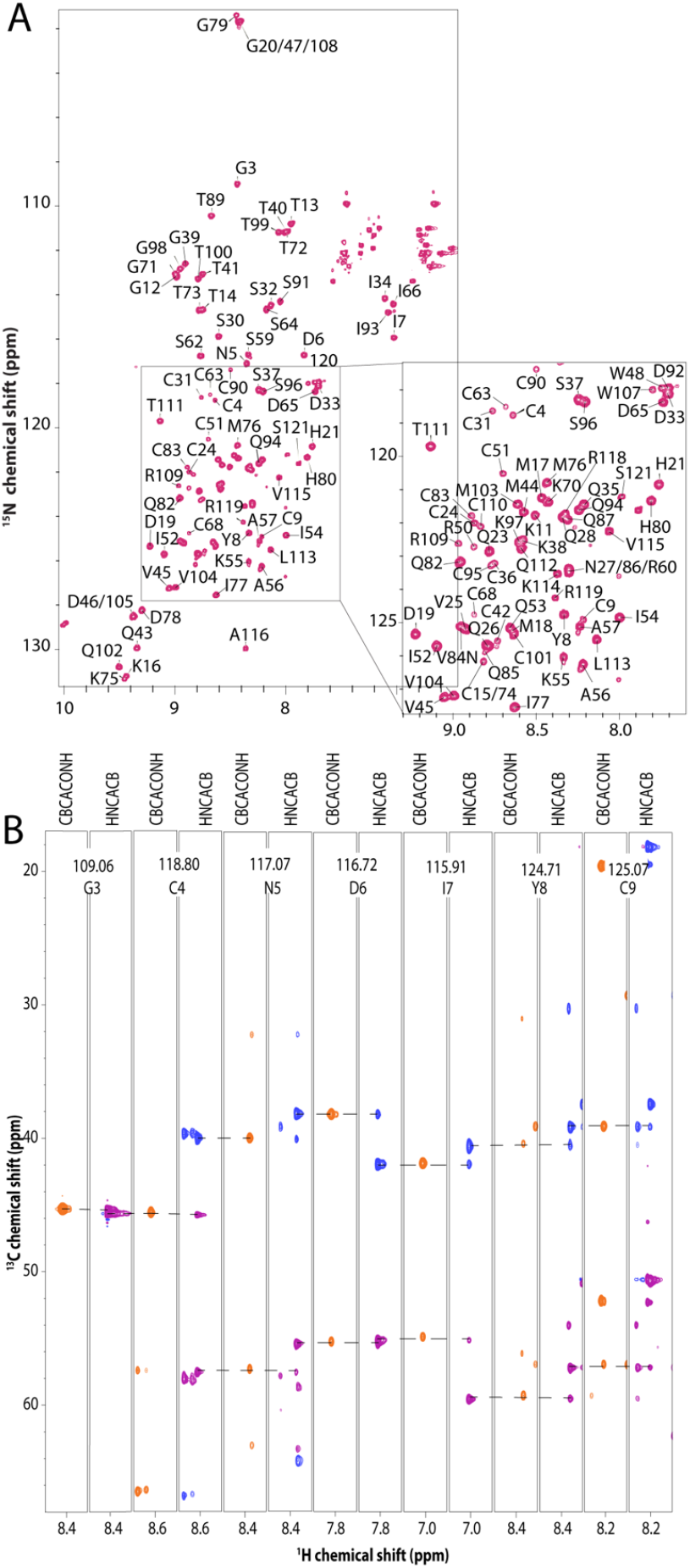
Solution NMR spectra of 4-mod CREMP. **Panel A**. 2D ^1^H-^15^N HSQC spectrum of 4-mod U-^15^N,^13^C-CREMP. The selected cross-peak assignments are shown. Unassigned peaks in the upper right corner correspond to side-chain amino groups of Gln and Asn residues. **Panel B**. Strips from 3D CBCA(CO)NH (black) and HNCACB (CA in magenta and CB in cyan) spectra, demonstrating the sequential backbone walk for residues G2-C9.

The high sensitivity and resolution of the HSQC spectrum, along with the presence of a sufficient number of unique residues in otherwise highly similar module sequences (Figure S3B), allow for nearly complete chemical shift assignments (Figure S6 and Table S2). ^15^N HSQC, HNCACB (Wittekind & Mueller, 1993), CBCA(CO)NH (Grzesiek & Bax, 1992), HNCA, HNCO (Powers et al., 1991), and HNCACO (Kay et al., 1990) were used for backbone assignments. CCONH (Grzesiek et al., 1993), HNHA (Vuister & Bax, 1993; 1994), HNHB (Grzesiek & Bax, 1992), HBHA(CO)NH (Grzesiek & Bax, 1993; Muhandiram & Kay, 1994), HCCONH (Grzesiek et al., 1993; Montelione et al., 1992), HBCBCGCDHD and HBCBCGCDCEHE (Yamazaki et al., 1993) were used for sidechain heteronuclear and proton assignments. Figure 3B shows the sequential assignments for Gly2-Cys9 in the first module, and Table S2 summarizes the assigned chemical shifts.

A ^15^N-edited NOESY (Marion et al., 1989a; Marion et al., 1989b; Zuiderweg & Fesik, 1989) spectrum with a 200 ms mixing time revealed a robust network of sequential, short, and medium-range NOE cross-peaks. Figure 4A displays some of these restraints for I7, D6, S59, and S62. Notably, comparatively few long-range contacts were detected in each module (Figure S7). Tables S3 and S4 summarize the details of intra- and inter-module correlations, respectively. Overall, we assigned 1448 distance restraints, including 335 medium-range (2 ≤ |i – j| ≤ 5) and 204 long-range (|i – j| ≥ 5) restraints (Table I).

**Figure 4.**
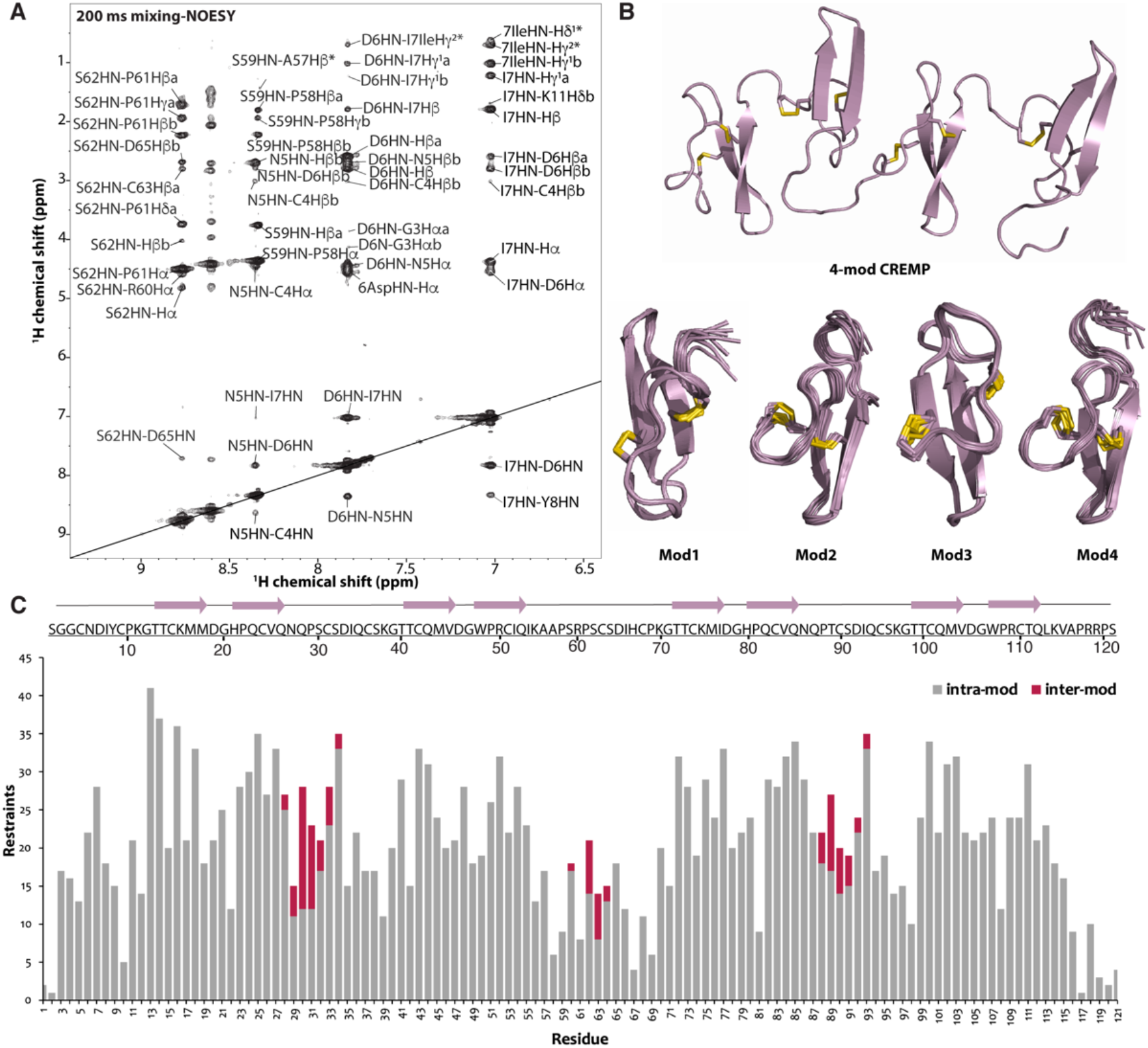
NOE assignments and a structural model of 4-mod CREMP. **Panel A**. A selected region of the ^15^N-edited NOESY spectrum of 4-module CREMP, highlighting the assigned NOE cross-peaks used for structure calculations. **Panel B**. On the top, the 4-mod CREMP model (cartoon representation) is shown. On the bottom, the 10 lowest-energy structural models calculated for each individual module (Mod1–Mod4) are displayed. Each module exhibits a well-defined fold with pairwise RMSD values of 4.9 ± 2.1 Å (backbone) and 5.3 ± 2.1 Å (heavy atom). **Panel C**. A bar graph showing the distribution of distance restraints used in the structure calculation. The total number of intra-module restraints (grey) and inter-module restraints (burgundy) plotted against the residue number. The primary sequence and secondary structure elements are shown on the top.

**Table I.**
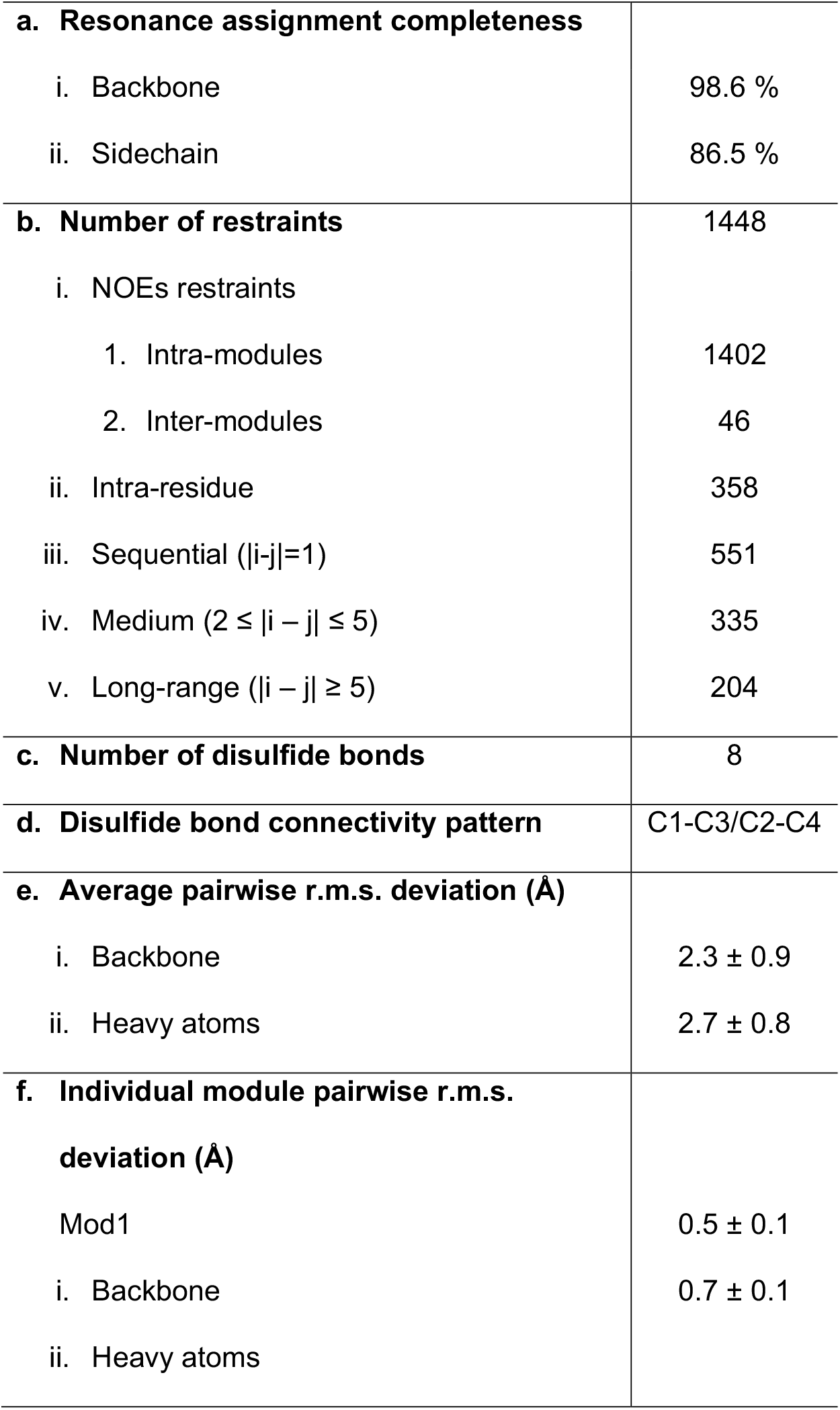
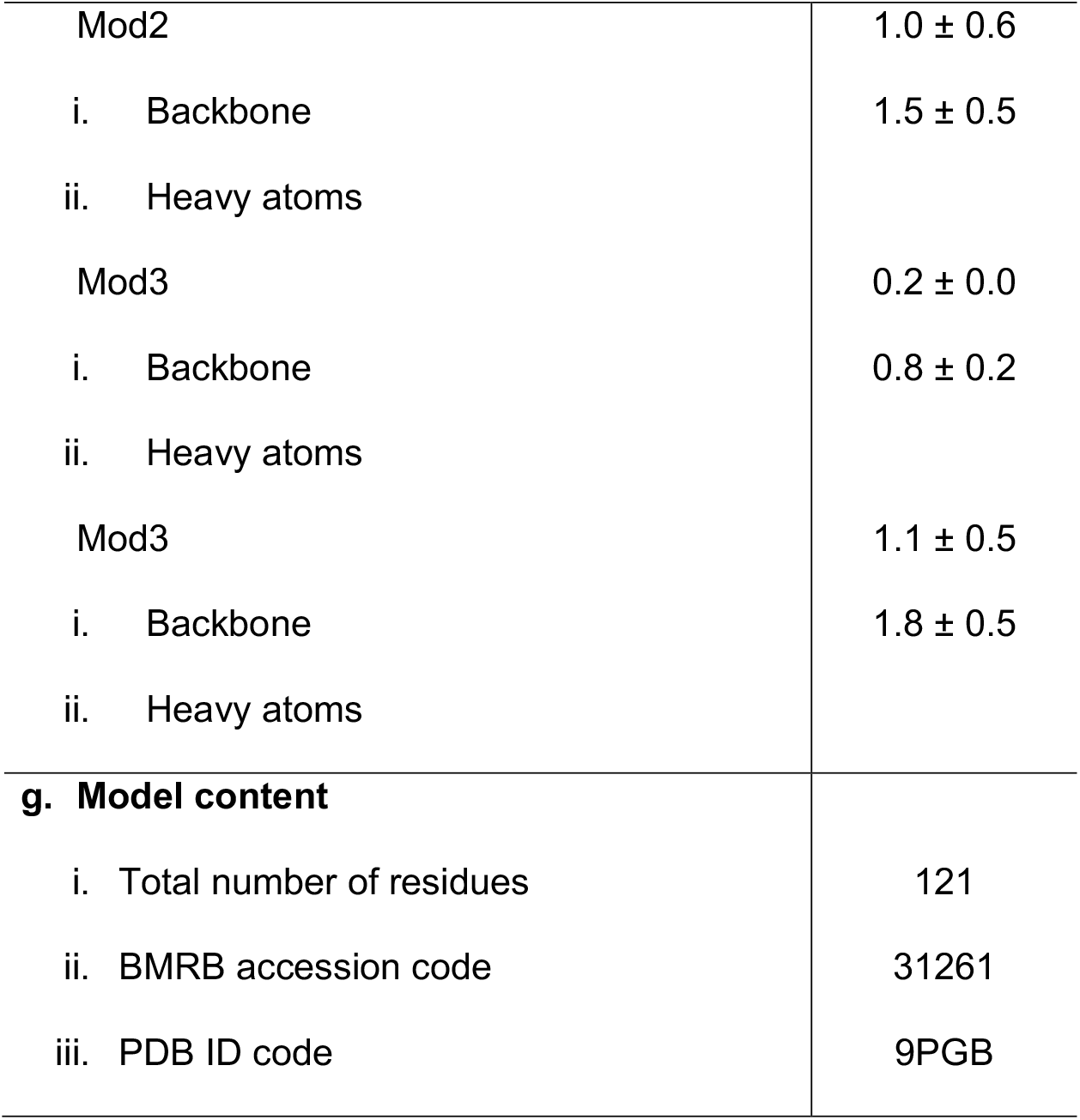
NMR and refinement statistics of 10 lowest-energy structures of 4-mod CREMP.

|  |  |
| --- | --- |
| **a. Resonance assignment completeness** |  |
| i. Backbone | 98.6 % |
| ii. Sidechain | 86.5 % |
| **b. Number of restraints** | 1448 |
| i. NOEs restraints |  |
| 1. Intra-modules | 1402 |
| 2. Inter-modules | 46 |
| ii. Intra-residue | 358 |
| iii. Sequential ($\vert i-j\vert =1$) | 551 |
| iv. Medium ($2 \leq \vert i-j\vert \leq 5$) | 335 |
| v. Long-range ($\vert i-j\vert \geq 5$) | 204 |
| **c. Number of disulfide bonds** | 8 |
| **d. Disulfide bond connectivity pattern** | C1-C3/C2-C4 |
| **e. Average pairwise r.m.s. deviation (Å)** |  |
| i. Backbone | $2.3 \pm 0.9$ |
| ii. Heavy atoms | $2.7 \pm 0.8$ |
| **f. Individual module pairwise r.m.s. deviation (Å)** |  |
| Mod1 | $0.5 \pm 0.1$ |
| i. Backbone | $0.7 \pm 0.1$ |
| ii. Heavy atoms |  |
| Mod2 | $1.0 \pm 0.6$ |
| i. Backbone | $1.5 \pm 0.5$ |
| ii. Heavy atoms |  |
| Mod3 | $0.2 \pm 0.0$ |
| i. Backbone | $0.8 \pm 0.2$ |
| ii. Heavy atoms |  |
| Mod3 | $1.1 \pm 0.5$ |
| i. Backbone | $1.8 \pm 0.5$ |
| ii. Heavy atoms |  |
| **g. Model content** |  |
| i. Total number of residues | 121 |
| ii. BMRB accession code | 31261 |
| iii. PDB ID code | 9PGB |

Our model for the 4-mod CREMP indicates an extended structure featuring four beta-hairpins separated by linker regions (Figure 4B and 4C). As shown in Figure 4D and Tables S3 and S4, there are a very small number of inter-module distance restraints (about 3%), and they are mainly sequential or medium range. In addition, there are fewer distance restraints in the loop regions connecting modules, especially in the modules with the longer linker (the second and fourth modules compared to the first and third ones). This result suggests that the protein adopts an extended, rather than compact, structure, with four modules being independently folded. The lack of inter-module NOEs leads to the structural heterogeneities and contributes to a large precision RMSD value, 2.3 ± 0.9 Å, in the overall model. In contrast, the precision RMSDs for the individual modules are 0.5 ± 0.1, 1.0 ± 0.6, 0.2 ± 0.0, and 1.1 ± 0.5 Å, as shown in the superposition of the 10 lowest-energy models for the individual modules (Figure 4B and Table I).

Notably, a comparison of our solution NMR structural model with the predicted AF3 structure (Abramson et al., 2024) shows a similar architecture (Figure S8). Both the NMR and AF3 models show an extended model with sparse inter-module contacts. Clear differences mainly arise from loops and linker regions connecting the beta-hairpins. These conformational differences are attributed to the inherent flexibility of loops/linkers in solution compared to the rigid X-ray structures, which served as the source for AF3 training (Fowler & Williamson, 2022; Jumper et al., 2021).

In addition to the notable conservation of sequence between C1 and C4 in Figure 2C, the inter-module regions also retain a considerable level of sequence conservation when linkers of the same length are compared. For illustration, the 20 modules in Figure 2B show an alternation of 6 and 11 amino acid linker lengths (between the last Cys of one module and the first of the next; Figure S9A and B). In the NMR structural model (Figure 4B), there is no clear explanation for this conservation of residues within either linkage type. It is possible that these regions may contribute to higher-order structures by facilitating critical intermolecular or intramolecular interactions within ESM fibers. Following the CREMP modules, the remaining 25% of the sequence contains 3 VWFD domains and 2 small cysteine-rich C8 domains (Figure 2A, with details in Figure S2B), and as noted earlier, these domains may provide additional interprotein disulfide linkages.

### Comparison of CREMP with selected disulfide-rich tandem repeat proteins

A brief consideration of selected tandem repeat structural proteins that are notably disulfide-rich offers a valuable perspective for the CREMP structural model established in this study. Many of these proteins contain epidermal growth factor (EGF)-like domains (Campbell & Bork, 1993). Although there is significant variability in the placement of Cys within three disulfide EGF modules (Tombling et al., 2020), the NOTCH-fifth domain (PDB 5FMA) exhibits the Cx_4_Cx_5_Cx_8_C spacing common to almost all CREMP modules and is compared with the third CREMP module in Figure S10. Both peptides display the same C1-C3 and C2-C4 connectivity and arrangement relative to the beta-hairpin structure, but the CREMP domain lacks the third disulfide in the C-terminal portion of EGF. Craik and colleagues have demonstrated that truncations of the terminal third of an EGF domain, which removes this final disulfide, can be both kinetically and thermodynamically stable (Tombling et al., 2020). One particularly intriguing example of an EGF repeat structural protein is found in octopus egg threads. Octovafibrin consists of 29 EGF domains, with all but one showing a spacing of 4, 5, and 8 amino acids between the first four cysteine residues (Wonderly et al., 2023).

The enormous disulfide-rich mosaic protein DUMPY from *Drosophila melanogaster* (22,949 amino acids; isoform Q) forms structural filaments between the fly epidermis and cuticle (Chu & Hayashi, 2021; Wilkin et al., 2000). DUMPY contains multiple domain types, including 308 EGF modules and 185 copies of a small 2-disulfide Dumpy module (DPY) (Wilkin et al., 2000). A comparison of the NMR structure of a DPY module with the third module of CREMP is presented in Figure 5A. Again, the DPY and CREMP modules show rather similar architecture, although they feature different consensus spacings (Cx_5_Cx_5_Cx_4_C for DPY and Cx_4_Cx_5_Cx_8_C for CREMP).

**Figure 5.**
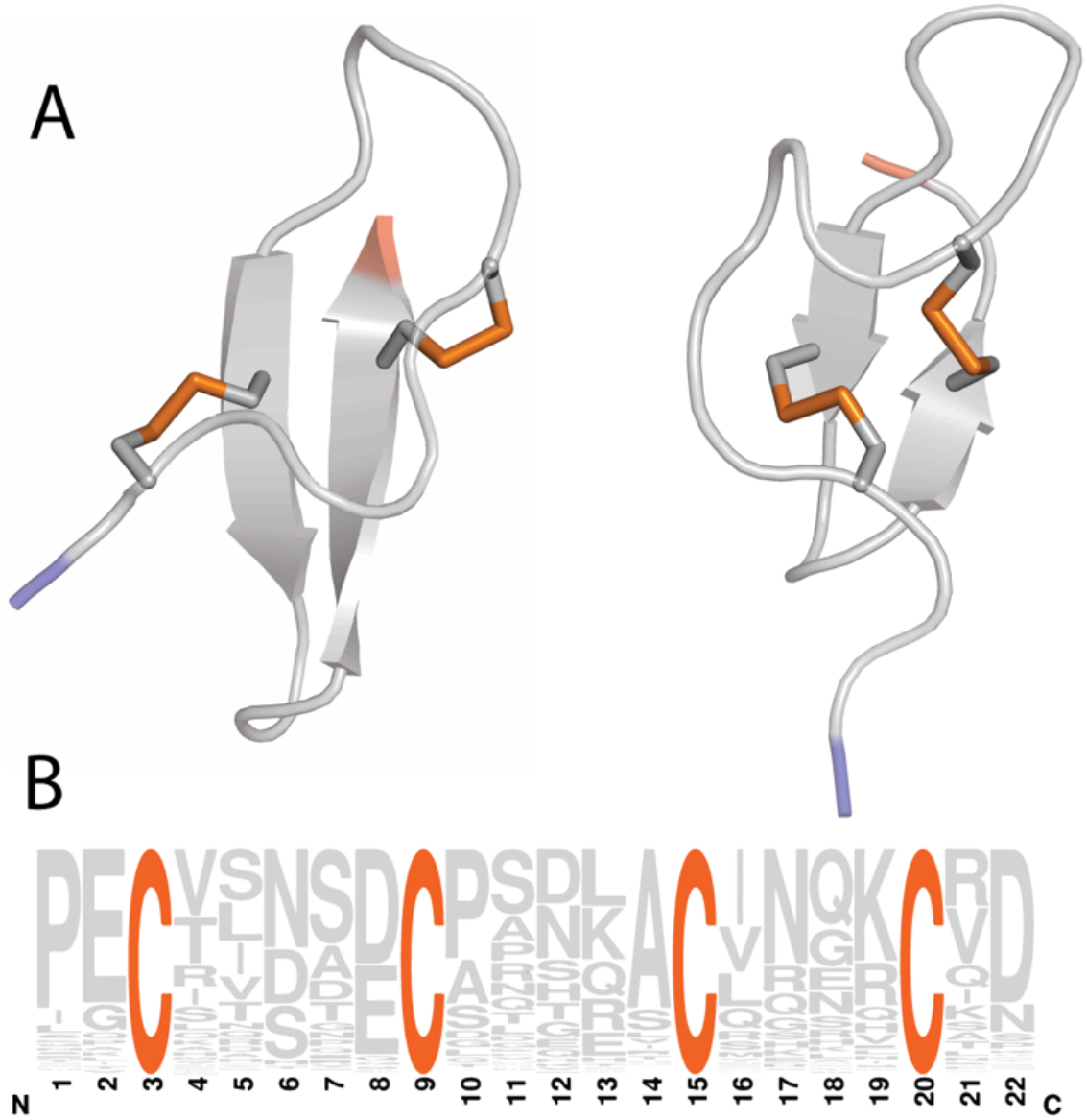
**Panel A**. A Comparison of the third CREMP module (left) with the structure of the DPY module (right; PDB 1OIG (Wilkin et al., 2000)). N- and C-termini are colored blue and red, respectively. **Panel B**. A WebLogo graphic of the sequence variation around the highly conserved 4 cysteine residues (spacing CX_5_CX_5_CX_4_C) in Dumpy Isoform Q.

Finally, a striking example of a disulfide-rich structural protein comes from the aquatic cocoons of the *Chironomus* midge (Case et al., 1997). This silk protein, Sp185 is 1698 amino acids long with a Cys content of 16.8%. Lacking an experimental structure, an AF3 model predicts that the 64 tandem repeats of a 2-disulfide module show the disulfide connectivity and overall resemblance to CREMP and DPY modules. A model of two successive modules is shown in Figure S11A, in which the last cysteine of each CxCxC motif is disulfide-bridged to the third cysteine residue in the next module. Unlike CREMP and DPY modules, these *Chironomus* modules vary markedly in length and cysteine spacings. Figure S11B illustrates this variability using a WebLogo representation of 19 amino acids following the first cysteine of all SP185 modules. Panel C compares the experimental structure of the 3rd CREMP module with module 46 of the silk protein predicted by AF3.

### Broader perspectives and future outlook

This study focuses on the structure of the CREMP modules from *Gallus gallus* and is not intended to survey the large number of CREMP sequences in the bird and reptile databases (Kodali et al., 2011). Although these structural proteins were first described in 2010 (Kodali et al., 2011), they are frequently annotated in the databases as zonadhesin-like, alpha-tectorin-like, Follistatin-like domain-containing protein, IgGFc-binding protein-like, and VWFD domain-containing protein. We note that all the avian and reptilian CREMP sequences surveyed to date show long runs of CREMP modules (see, for example, Figures 2A and S2) and hence would appear to warrant a unified descriptor. The remainder of the chicken CREMP sequence features three VWFD domains and two cysteine-rich C8 domains. VWFD domains are involved in multimerization of von Willebrand factor and mucins to create extremely long linear functional assemblies (Javitt et al., 2020; Perez-Vilar & Hill, 1999; Thornton et al., 2018). The roles of VWF domains in CREMP structure and function, and the possible existence of disulfide-linked multimers in the disulfide-rich avian ESM remain to be investigated. It should be noted that there are negligible free reduced Cys residues in chicken ESM (Raje, 2003).

All CREMP sequences surveyed in this study feature multiple tandem repeats of a small 2-disulfide module with almost invariant Cx_4_Cx_5_Cx_8_C spacing (Kodali et al., 2011). It seems likely that CREMP protein serves as an integral structural component of the fibrous structures of the avian ESM. Indeed, CREMP appears to be abundant in the chicken ESM and has the highest Cys content of all candidate proteins within that structure (Ahmed et al., 2019). We therefore examined cross sections of unstained ESM using energy-filtered transmission electron microscopy edited for sulfur abundance (Egerton, 2011). The proteinaceous fibers show a core that is notably more sulfur-rich than the mantle (Figure S12). Recognizing that the average cysteine and methionine content of ESM proteins is approximately 10.4% and 3.4%, respectively (Kodali et al., 2011), the elevated sulfur level in the core is at least consistent with the presence of CREMP at that location. Further studies are needed to deconstruct ESM fibers, allowing for the reliable quantification of the protein complement in both the core and mantle. Finally, since small disulfide-rich peptides have been reported as vehicles for the generation of surfaces that confer opportunities for drug and material design (Tombling et al., 2020), the robustness, multivalency, and tolerance to a range of linker sizes inherent in CREMP modules may provide an extended scaffold for designed functional polypeptides.

## Materials and Methods

### Materials

Common chemicals were purchased from Sigma-Aldrich and Fisher Scientific. The new construct (cloned into pMAL-c5X) was ordered from GenScript. The SHuffle (T7) *Escherichia coli* competent cell line used for protein expression was purchased from New England Biolabs, Inc. ^15^NH_4_Cl, D_2_O, and U-^13^C_6_-glucose were purchased from Cambridge Laboratories, Inc. Amylose resin was from New England Biolabs. Cation-exchange chromatography columns (Hitrap SP HP column) were purchased from Cytiva.

### Plasmid design

The sequence of 4-mod CREMP was initially obtained from a normalized chicken reproductive tract cDNA library (Sequence ID: BM439825.1) as described earlier (Kodali et al., 2011). In the present work, the same sequence for the CREMP modules was codon-optimized for expression in *E. coli* and was subcloned into the pMAL-c5X vector (Figure S3). The previous construct (Kodali et al., 2011) contained an N-terminal hexa-HIS tag, subsequently removed via TEV protease. The present construct includes a maltose-binding protein (MBP) fusion partner upstream of the 4-mod CREMP sequence (Figure S3). Additionally, we inserted a TEV cleavage site (ENLYFQ|S) followed by an intervening GG dipeptide immediately before the 4-mod CREMP sequence so that the released CREMP fragment would be the same as that expressed previously by Kodali et al. (Kodali et al., 2011). TEV cleavage then allowed for the purification of pure 4-mod CREMP using a single cation-exchange step (Figure S4). This newer procedure avoids a persistent contaminant encountered during the earlier purification of the His-tagged material (Kodali et al., 2011).

### Protein expression and isotope labeling

The plasmid construct was transformed into SHuffle (T7) *E. coli* cells (Lobstein et al., 2012). For protein expression, 2 L of LB media were inoculated using an overnight starter culture and incubated at 30 °C while shaking at 180 rpm until an OD_600_ of ∼ 0.8-1.0 was reached. The cells were harvested by centrifugation at 5,000xg for 15 minutes at 4 °C and washed using M9 minimal salts buffer. The washed cells were then resuspended in 0.5 L of isotopically-labeled M9 media containing 0.5 g ^15^NH_4_Cl and 2 g U-^13^C_6_-glucose and shaken for 1 h at 16 °C for growth recovery (Marley et al., 2001). Cells were induced overnight using 0.5 mM IPTG at 16 °C. The cells were then pelleted by centrifugation at 5,000×g for 15 min, resuspended in 30 mL lysis buffer (20 mM Tris-HCl, 200 mM NaCl, 1 mM EDTA, pH 7.4), and stored at -80 °C before purification.

### Protein purification

Cell disruption was performed on ice using two passages through a manually operated French pressure cell at 10,000 to 15,000 psi. The suspension of broken cells was then subject to 5 min intermittent sonication (15 s pulse with 45 s cooling on ice) followed by centrifugation for one hour at 12,000×g, 4 °C. The clear supernatant was loaded by gravity onto a 40 mL amylose resin column and washed with 10 column volumes of buffer (20 mM Tris-HCl, 200 mM NaCl, 1 mM EDTA, pH 7.4). The MBP-CREMP construct was then eluted, including 10 mM maltose in the same buffer. Fractions were analyzed by SDS-PAGE, and those containing MBP-CREMP were pooled (in aggregate about 80 mL) and then dialyzed for 3 h against 25 mM potassium phosphate buffer (KPi), pH 7.75 at 4 °C. This solution was then supplemented with TEV protease (1% w/w), and the mixture was incubated overnight at 4 °C. The solution was then dialyzed against 25 mM KPi buffer, pH 6.8 at 4° C, followed by centrifuging for 15 min at 12,000×g, 4 °C. The TEV-treated clear solution was then subjected to cation-exchange chromatography on a 5 mL column yielding baseline separation of MBP, CREMP, and TEV. The total yield of purified protein from the initial 2L culture was estimated to be 10 mg using an extinction coefficient of 13,490 M^-1^cm^-1^ at 280 nm (predicted by ProtParam (Gasteiger et al., 2005)). The purity of the samples and their approximate molecular weight were evaluated using SDS-PAGE (Figure S4A). Mass spectrometry (on a Waters Xevo G2-S QT) gave a molecular weight of 13,094 Da, consistent with the value expected for the TEV-cleaved oxidized product (Figure S4B).

### Crystallization trials

The TEV-cleaved 4-mod CREMP and its MBP-fused construct were subjected to crystallization trials, both in-house and in 1536-well screening experiments at the Hauptman Wood Institute National Crystallization Center (Buffalo, New York) (Lynch et al., 2023).

### Sample preparation for solution NMR

Collectively, four NMR samples were prepared by dissolving isotopically labeled 4-mod CREMP (∼ 400 µM, ∼ 5mg/mL) in various ratios of H_2_O/D_2_O containing 25 mM KPi, 1 mM EDTA, pH 6.2. U-^15^N-CREMP and U-^13^C,^15^N-CREMP were dissolved in the above buffer using 90% H_2_O/10% D_2_O for performing 2D and 3D NMR experiments. ^15^N-edited NOESY-HSQC experiments were conducted on U-^13^C,^15^N-CREMP dissolved in 95% H_2_O/ 5% D_2_O. For recording ^13^C-edited NOESY-HSQC spectra, U-^13^C,^15^N-CREMP was dissolved in 99.9% D_2_O, and the pH of the sample was adjusted to 5.8. Lastly, a U-^15^N-4-mod CREMP with ^14^N-Trp (Figure S5) was dissolved in the buffer containing 95% H_2_O/ 5% D_2_O for unambiguous assignments of two tryptophan residues (W48 and W107).

### Solution NMR data acquisition, processing, and analysis

All NMR spectra were recorded at 298 K on a 14.1 T (600 MHz) Bruker NEO spectrometer equipped with a CryoProbe. Larmor frequencies were 600.13, 150.91, and 60.82 MHz for ^1^H, ^13^C, and ^15^N, respectively. The chemical shifts were referenced indirectly to 2,2-dimethyl-2-silapentane-5-sulfonate. Non-uniform sampling was used for all experiments. We recorded a total of 16 2D and 3D spectra for backbone, side chain, and distance restraints assignments. Sequential assignments of the backbone resonances were achieved by 2D and 3D-heteronuclear experiments (^15^N-HSQC, HNCACB (Wittekind & Mueller, 1993), CBCA(CO)NH (Grzesiek & Bax, 1992), HNCO (Powers et al., 1991), HNCA, and HN(CA)CO (Kay et al., 1990)). The 3D-CCCONH experiment was acquired for carbon side chain assignments (Grzesiek et al., 1993). ^15^N-edited NOESY spectra with a mixing time of 200 ms were acquired for determining H-based distance restraints. Table S1 details the acquisition and processing parameters.

NMR spectra were processed in NMRpipe using the SMILE reconstruction protocol (Ying et al., 2017) under the NMRBox environment (Maciejewski et al., 2017). Chemical shift assignments were performed in CCPN 2.4.2. (Skinner et al., 2016) DANGLE (Dihedral ANgles from Global likelihood Estimates) was employed for dihedral restraints calculation (φ and ψ) and secondary structure prediction (Skinner et al., 2016). AlphaFold3 was used for predicting the initial 4-mod CREMP structural model (Jumper et al., 2021). The well-folded parts of this model were then used as input into CCPN 2.4.2 to derive a synthetic NOE list. Comparison of this list with ^15^N-edited NOESY-HSQC spectra facilitated the assignment of intra-residue, short, medium, and long-range NOE peaks (Li et al., 2023; Tejero et al., 2022). Xplor-NIH version 2.51 (Schwieters et al., 2018; Schwieters et al., 2006; Schwieters et al., 2003) and PYMOL (The PyMol Molecular Graphics System, Version 2.5.2 Schrödinger, LLC) were used for structure calculations and visualization, respectively.

### Structure calculation

The structure calculation protocol employed here closely followed the approach we developed previously (Lu et al., 2020a; Russell et al., 2019). Structure calculations were performed using Xplor-NIH version 2.5.1. Standard terms for bond lengths, bond angles, and improper angles were utilized to enforce correct covalent geometry. A statistical torsion angle potential (Bermejo et al., 2012) and the gyration volume term were implemented (Schwieters & Clore, 2008). A hydrogen bond database term, HBPot, was used to enhance hydrogen bond geometries (Schwieters et al., 2020). Backbone dihedral restraints in the form of φ/ψ angles were predicted using DANGLE from the experimental solution NMR ^13^C and ^15^N chemical shifts.

Folding calculations were seeded from primary sequence extended strands, and disulfide bonds were explicitly defined. One thousand structures were calculated using molecular dynamics simulated annealing in torsion angle space with two successive annealing schedules and a final gradient minimization in Cartesian space. The structure calculation began with a 3,500 K constant-temperature molecular dynamics run for the shorter of 800 ps or 8,000 steps with the time step size allowed to float to maintain constant energy, within a tolerance. The initial velocities were randomized about a Maxwell distribution using a starting temperature of 3,500 K. Following this initial dynamics calculation, a simulated annealing calculation was performed where the temperature was reduced to 100 K in steps of 25 K.

At each temperature, dynamics was run for the shorter of 0.4 ps or 200 steps. Force constants for distance restraints were ramped from 10 to 50 kcal mol^−1^ Å^−2^. The dihedral angle restraints were disabled for high-temperature dynamics at 3,500 K, but enabled during simulated annealing with a force constant of 200 kcal mol^−1^ rad^−2^. The gyration volume force constant was geometrically scaled from 0.002 to 1. After simulated annealing, the structures were minimized using a Powell energy minimization scheme.

Subsequently, the ten lowest-energy structures from the fold calculations were selected as input for refinement calculations. As with the folding calculations, one thousand structures were generated. Simulated annealing at 3,000 K was performed for 10 ps or 5,000 steps, whichever was completed first. The starting time step was 1 fs and was self-adjusted in subsequent steps to ensure conservation of energy. The initial velocities were randomized about a Maxwell distribution using the starting temperature of 3,000 K. The temperature was subsequently reduced to 25 K in steps of 12.5 K. At each temperature, the initial default time step was 1 fs, and a 0.2-ps dynamics run was performed. Force constants for distance restraints were ramped from 2 to 30 kcal mol^−1^ Å^−2^. The dihedral restraint force constants were set to 10 kcal mol^−1^ rad^−2^ for high-temperature dynamics at 3,000 K and 200 kcal mol^−1^ rad^−2^ during cooling. The gyration volume force constant was scaled from 0.002 to 1. The annealed structures were minimized using a Powell energy minimization scheme. Of the one thousand structures generated, the ten lowest-energy structures from each run were selected for the next step. This was repeated subsequently to total four iterations of refinement.

### Preparation of Eggshell Membrane

Large white eggs were obtained from a local supermarket, cracked at their widest point and emptied before washing their insides for 5 min under a rapid stream of first tap and then distilled water. The blunt-end halves, with their resident air sac, were discarded. The pointed-end halves were filled with 100 mM EDTA, pH 7, and incubated for 6 hours at room temperature. The halves were then emptied and immersed in distilled water in a large crystallizing dish to allow facile detachment of the eggshell membrane. Membrane pieces were subsequently cut under water and transferred to a small volume of 1 mM EDTA for storage at 4 °C prior to examination by SEM and TEM.

### Serial block-face electron microscopy

For serial block-face processing, eggshell membranes were fixed overnight in 2% paraformaldehyde and 2% glutaraldehyde in 0.1 M sodium cacodylate buffer, pH 7.4. The sample was processed according to a modified protocol developed by Hua et al. (Hua et al., 2015) that included serial incubation steps with solutions of osmium tetroxide, potassium ferrocyanide, thiocarbohydrazide, uranyl acetate, and Walton’s lead aspartate.(Walton, 1979) The ESM sample was dehydrated with an ascending ethanol series (50, 75, 95, and 100% v/v), then infiltrated with Embed 812 resin (hard formulation; Electron Microscopy Sciences) and polymerized for 24 h in a 60°C oven. The embedded sample was then mounted on specimen stubs using Epo-Tek H20E conductive epoxy (Ted Pella Inc.), trimmed using an ultramicrotome (Leica EM UC7), and sputter coated with 12.5 nm of platinum (Leica EM ACE600). The SBF image stacks were acquired using a ThermoFisher Scientific Apreo Volumescope SEM using MAPS software in low vacuum mode at 50 Pa with an accelerating voltage of 2.5 keV and at a beam current of 0.1 nA. The smaller image stack had a 32.77 µm x 32.77 μm in- plane field of view with an 8 nm/pixel resolution, and the larger image stack had a 98.3 µm x 98.3 µm in-plane field of view with a 12 nm/pixel resolution. A total of 600 slices were collected with 50 nm spacing, resulting in an image stack along a 30 μm total axial length. Image processing, rendering, and video animations were conducted in Amira 3D 2022 software (ThermoFisher Scientific).

### Energy-filtered Transmission Electron Microscopy

Eggshell membrane pieces were fixed overnight in 2% glutaraldehyde and 2% paraformaldehyde in 0.1 M sodium cacodylate buffer pH 7.4. The membranes were washed with 0.1 M sodium cacodylate buffer pH 7.4, dehydrated in an acetone series, infiltrated gradually with Embed-812 resin, and polymerized at 60 °C overnight. Ultrathin (50-60 nm) sections were cut using a Leica UC7 ultramicrotome and collected onto 400-mesh copper grids with a carbon film. A Zeiss Libra 120 transmission electron microscope with an in-column Omega-type energy filter and a Gatan Ultrascan 1000 CCD was used to characterize sulfur in the sections. Data were collected at 120 kV using a Gatan 626 cryo-holder to keep the sections at -165 to - 170 °C. Sulfur maps were created using a 3-window method (Egerton, 2011) with a post-edge energy of 171 eV, a pre-edge1 energy of 127 eV, a pre-edge2 energy of 149 eV, and a slit width of 20 eV. The background model used for mapping was a smoothed R-power law.

## SUPPLEMENTARY MATERIAL

(File name: CREMP_Supplementary): contains Supplementary Figures S1-S12 and Tables S1-S4, together with legends and associated references.

## SUPPLEMENTARY VIDEOS

Video ESM S1 and Video ESM S2 provide a 3D. representation of a limited (S1) and more extensive (S2) section of chicken eggshell membrane visualized using serial block-face scanning electron microscopy.

## ACKNOWLEDGMENTS

We thank Vamsi Kodali for recognizing the CREMP protein family and for continuing advice. We are grateful to Sam Scinto for help with plasmid design and Karl Schmitz and Monika Prorok for help with initial crystal screening. We thank Shi Bai for assistance with NMR instrument setup, and Andrew P Hinck for providing sample experimental settings and processing scripts for NMR experiments.

## AUTHOR CONTRIBUTIONS

C.T. and T.P. conceived the project and guided the work. S.Z. prepared the NMR samples and performed the NMR data analysis. T.P. performed the NMR experiments and processed the data. S.Z. analyzed and interpreted the spectra. R.W.R. wrote scripts for structure calculations. R.W.R and S.Z. analyzed the calculation results and structure visualizations. J.R. and S.M. conducted the EM measurements on ESM, and J.C. visualized and interpreted the SBF-SEM images. C.T., S.Z., and T.P. took the lead in writing the manuscript. All authors discussed the results and contributed to the manuscript preparation.

## DATA AVAILABILITY

Coordinates of the solution NMR structure have been deposited at the Protein Data Bank with accession code PDB ID 9PGB. The solution NMR chemical shifts are deposited at the Biological Magnetic Resonance Bank (BMRB) under BMRB entry ID 31261. The electron microscopy data in this study are openly available in EMBL-EBI at https://doi.org/10.6019/S-BSST2148, accession number S-BSST2148.

## Abbreviations

AF3: AlphaFold3
C8: Cysteine-Rich C8 Domain
CcpNMR: Collaborative Computing Project for NMR
CREMP: Cysteine-Rich Eggshell Membrane Protein
DANGLE: Dihedral ANgles from Global Likelihood Estimates
DPY: Dumpy Protein Module
EGF: Epidermal Growth Factor
HBPot: Hydrogen Bond Potential
HSQC: Heteronuclear Single Quantum Coherence
IPTG: Isopropyl β-D-1-thiogalactopyranoside
MBP: Maltose Binding Protein
NEB: New England Biolabs
NEO: Bruker NEO spectrometer
NOE: Nuclear Overhauser Effect
NOESY: Nuclear Overhauser Effect Spectroscopy
NOTCH: Neurogenic locus notch homolog
NUS: Non-Uniform Sampling
SBF-SEM: Serial block-face scanning electron microscopy
SMILE: Spectral Processing Method (Sparse Multidimensional Iterative Lineshape-Enhanced)
TEV: Tobacco Etch Virus
VWFD: von Willebrand Factor type D domain

