## Supplementary Figures and Tables for "Structure of the Disulfide-rich Modules of a Striking Tandem Repeat Protein, Avian Cysteine-Rich Eggshell Membrane Protein": CREMP_Supplementary material.pdf

**Table of Contents**

**Figure S1 |** Serial block-face electron microscope image of a more extended view of the eggshell membrane ........ S3

**Figure S2 |** Amino Acid Sequence of Chicken CREMP ........ S4

**Figure S3 |** Plasmid Map of pMAL-c5X-CREMP Construct. ........ S5

**Figure S4 |** Validation of TEV-cleaved 4-Module CREMP sample purity and molecular weight ........ S5

**Figure S5 |** $^{1}H$-$^{15}N$ HSQC spectra of 4-mod CREMP highlighting tryptophan resonances ........ S6

**Figure S6 |** Graphical summary of chemical shift assignments for U-$^{13}C$,$^{15}N$ 4-mod CREMP ........ S6

**Figure S7 |** Distribution of the medium and long-range distance restraints inside the individual modules ........ S7

**Figure S8 |** Structural comparison of solution NMR (left) and AF3 (Abramson et al., 2024) (right) models of the third CREMP module ........ S7

**Figure S9 |** Weblogo comparison of the short and long linker regions from the sequence shown in Figure 2B ........ S7

**Figure S10 |** Structural comparison of the third module of 4-mod CREMP (left) with an X-ray structure of EGF NOTCH domain (right, residues 250-294 of human Notch homolog protein 4; PDB 5fma) ........ S8

**Figure S11 |** Chironomus midge silk SP185 ........ S8

**Figure S12 |** Electron Energy-Loss Spectroscopy (A), brightfield (B), and Energy-filtered TEM (C) images of eggshell membrane ........ S9

**Table SI |** Summary of solution NMR experiments*. ........ S10

**Table SII |** NMR chemical shifts for 4-mod CREMP ........ S11

**Table SII (con'd) |** NMR chemical shifts for 4-mod CREMP ........ S16

**Table SIII |** Intra-module distance restraints ........ S21

**Table SIV |** Inter-module distance restraints ........ S21

**REFERENCES** ........ S21

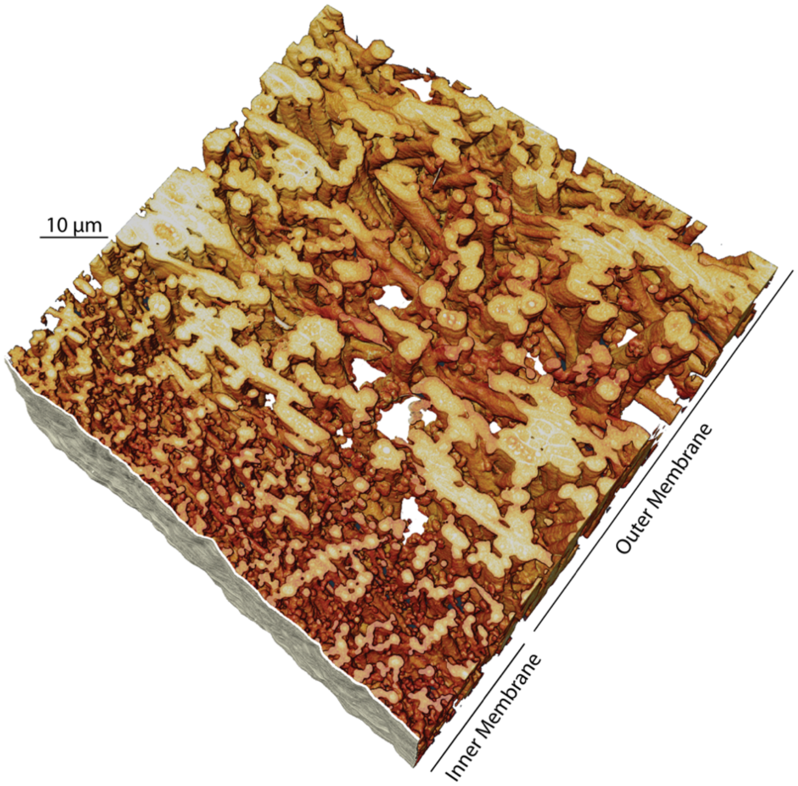

**Figure S1 | Serial block-face electron microscope image of a more extended view of eggshell membrane.** ESM was fixed, stained, and imaged as before (Figure 1). An accompanying video (ESM video S2) provides more details.

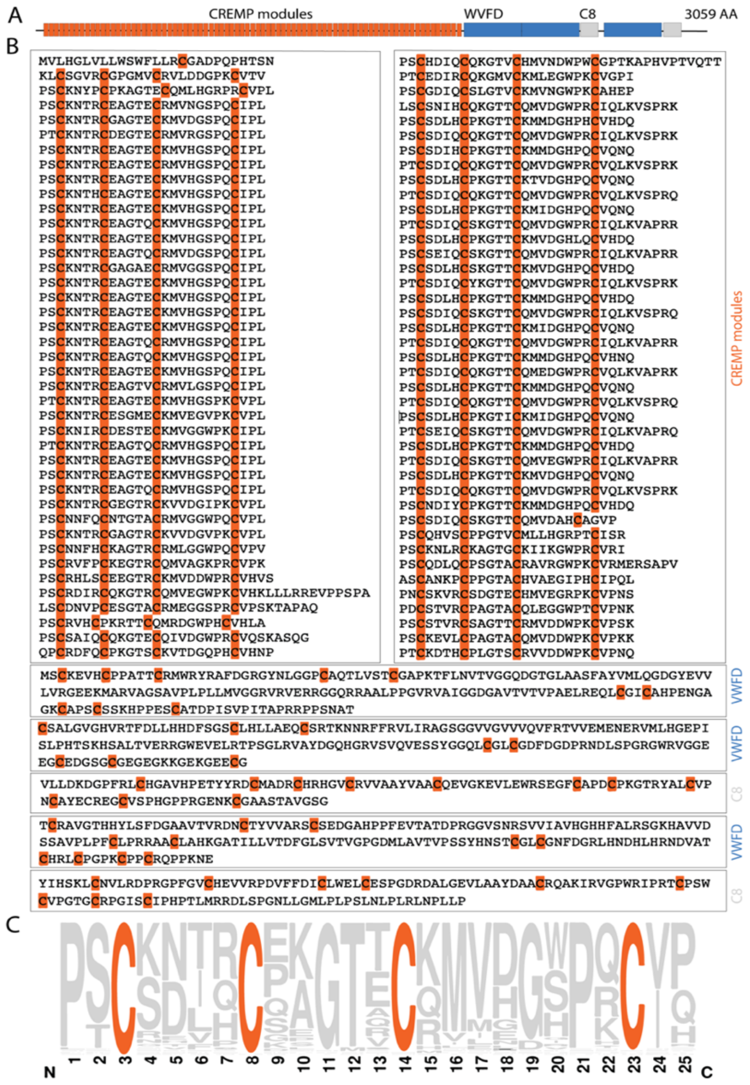

**Figure S2 | Amino Acid Sequence of Chicken CREMP (**Zonadhesin isoform X2; RefSeq: XP_040551000.1). **Panel A**. Schematic domain organization. **Panel B**. Full sequence segregating CREMP modules by line (arbitrarily starting here two residues before the first cysteine residue of each 4 Cys module; see main text). Spacings between C4 of one module and C1 of the next are often 5, 6, or 11 amino acid residues. The three von Willebrand Factor Type D Domains (VWFD) and the two C8 domains (PROSITE (Sigrist et al., 2013) and the NCBI Conserved Domains database (Lu et al., 2020)) reside at the C-terminus of this 3059 residue protein. **Panel C**. A WebLogo representation of all CREMP modules illustrates the striking conservation of cysteine spacing within this tandem repeat protein.

A

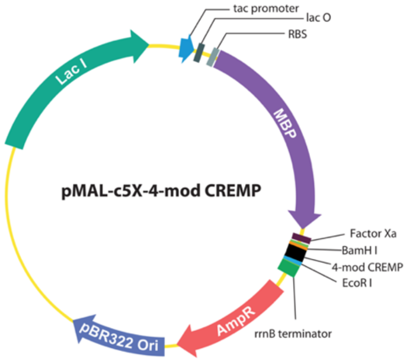

B

```
ENLYFQ|SGG
CNDIYCPKGTTCKMMDGHPQCVQNQPS
CSDIQCSKGTTCQMVDGWPRCIQIKAAPSRPS
CSDIHCPKGTTCKMIDGHPQCVQNQPT
CSDIQCSKGTTCQMVDGWPRCTQLKVAPRRPS
```

**Figure S3 | Plasmid Map of pMAL-c5X-CREMP Construct. A**. The plasmid map (adapted from GenScript) shows the pMAL-c5X vector with the 4-mod CREMP sequence inserted between the BamHI and EcoRI restriction sites. The construct includes a maltose-binding protein (MBP) tag for expression and purification, followed by a TEV protease cleavage site and the 4-mod CREMP sequence (shown in **B**). Key features such as the origin of replication (ori), antibiotic resistance marker (AmpR), and promoter region (Ptac) are indicated.

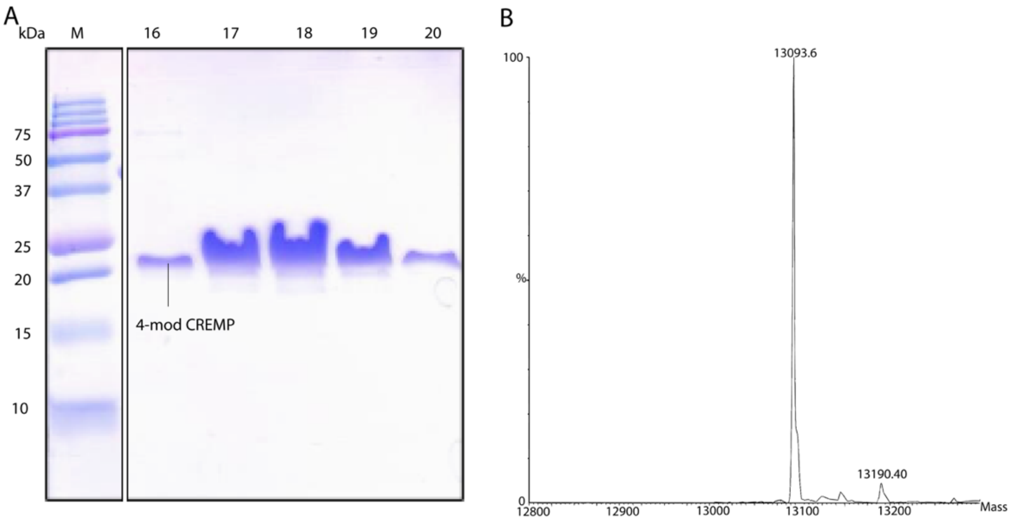

**Figure S4 | Validation of TEV-cleaved 4-Module CREMP sample purity and molecular weight. Panel A**. SDS-PAGE analysis of purified 4-mod CREMP. The Purification of TEV-cleaved 4-mod CREMP was performed using cation exchange chromatography (HP SP column). Eluted fractions 16-20 were collected for NMR experiments. While the actual mass of 4-mod CREMP is 13110 in its reduced form, it appears in its oxidized form on SDS PAGE as a single band at about 20 kDa, reflecting the presumed extended conformation of the 8-disulfide construct (Kodali et al., 2011). The marker lane is shown at the left. **Panel B**. Mass spectrometry analysis of oxidized 4-mod CREMP. The observed mass of 13094 Da corresponds to the theoretical mass of 4-mod CREMP with a full complement of 8 disulfide bonds.

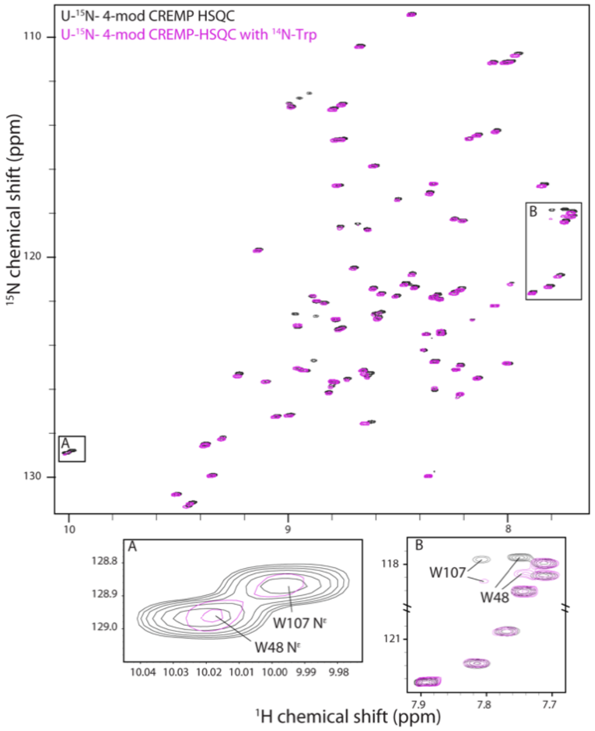

**Figure S5 | $^{1}H$-$^{15}N$ HSQC spectra of 4-mod CREMP highlighting tryptophan resonances.** The $^{1}H$-$^{15}N$ HSQC spectrum of 4-mod CREMP expressed in $^{15}N$-labeled M9 media supplemented with 50 mg/L of $^{14}N$-Trp (magenta) is overlaid with the spectrum of U-$^{15}N$-labeled 4-mod CREMP (black). The incorporation of $^{14}N$-Trp was performed to suppress the Trp sidechain (Ne) (inset A) and backbone amide (inset B) signals in the magenta spectrum.

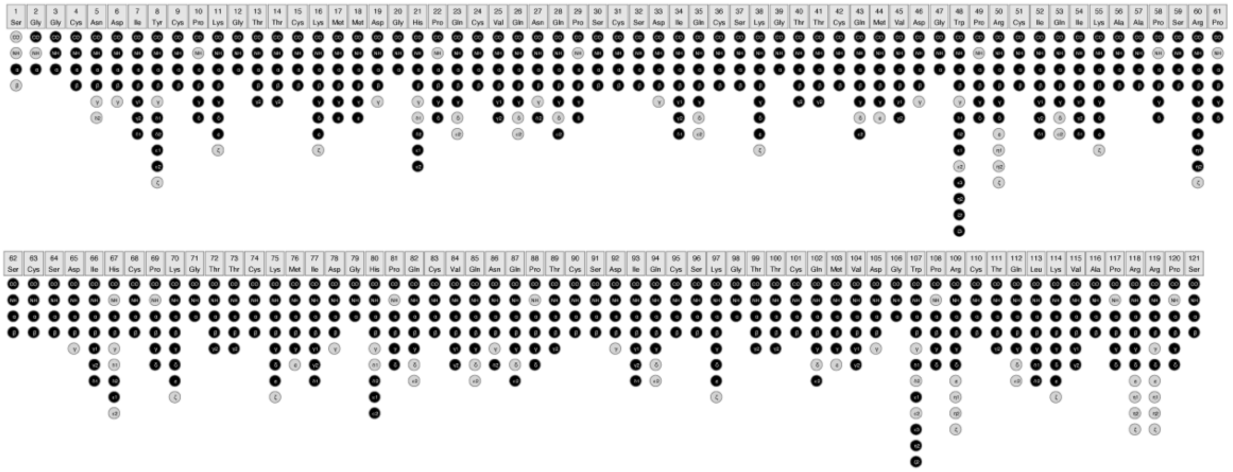

**Figure S6 | Graphical summary of chemical shift assignments for U-$^{13}C$,$^{15}N$ 4-mod CREMP.** Assigned atoms are depicted in black, while unassigned atoms are shown in grey. The summary highlights the completeness of the resonance assignments.

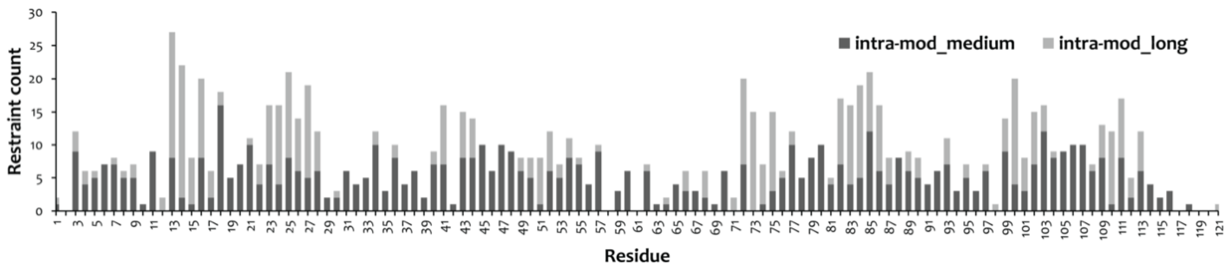

**Figure S7 | Distribution of the medium and long-range distance restraints inside the individual modules.** The bar graph illustrates the distribution of medium (dark grey) and long-range (light grey) distance restraints by residue number.

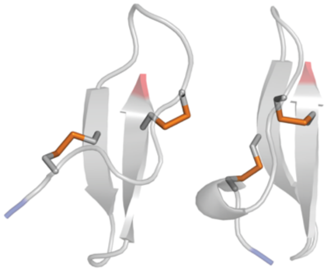

**Figure S8 | Structural comparison of solution NMR (left) and AF3 (Abramson et al., 2024) (right) models of the third CREMP module.** N- and C-termini are colored blue and red, respectively.

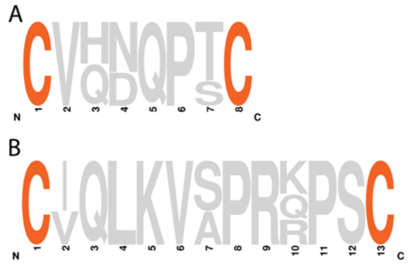

**Figure S9 | Weblogo comparison of the short and long linker regions from the sequence shown in Figure 2B.** Position 1 in panels A and B reflect the 4th Cys residue in one module with either a 6 or 11 amino acid linker region before the first cysteine of the next module.

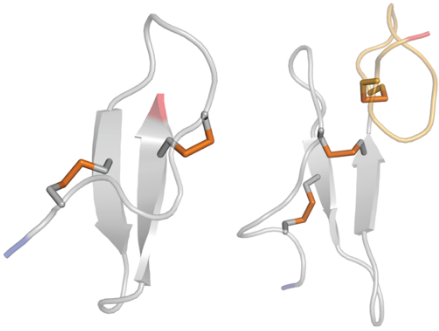

**Figure S10 | Structural comparison of the third module of 4-mod CREMP (left) with an X-ray structure of EGF NOTCH domain (right, residues 250-294 of human Notch homolog protein 4; PDB 5fma)**. The N-terminal region of the EGF module contains C1-C3 and C2-C4 disulfides (grey), with the C-terminal third (light orange) contains the C5-C6 disulfide absent in CREMP modules.

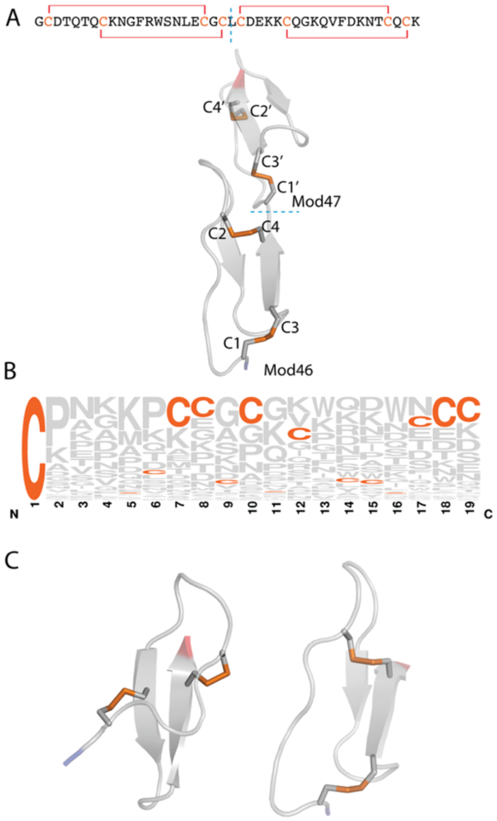

**Figure S11 | Chironomus midge silk SP185. Panel A**. Shows a high-confidence AF3 model of modules 46 and 47 selected from the 64 modules linked by CxCxC motifs. The corresponding amino acid sequence is shown, depicting the C1-C3 and C2-C4 disulfides seen in the structure. A model of the entire SP185 structure shows that all 64 modules apparently show the same disulfide connectivity and overall architecture (not shown). Nevertheless, there are wide variations in the placement of the 4 Cys residues and the length and sequence of the intervening regions. **Panel B**. Illustrates this variability using a WebLogo representation of 19 amino acids following the first cysteine of all SP185 modules. **Panel C**. Shows a comparison of the third module of the 4-mod CREMP (left) with module 46 of the silk SP185 (right).

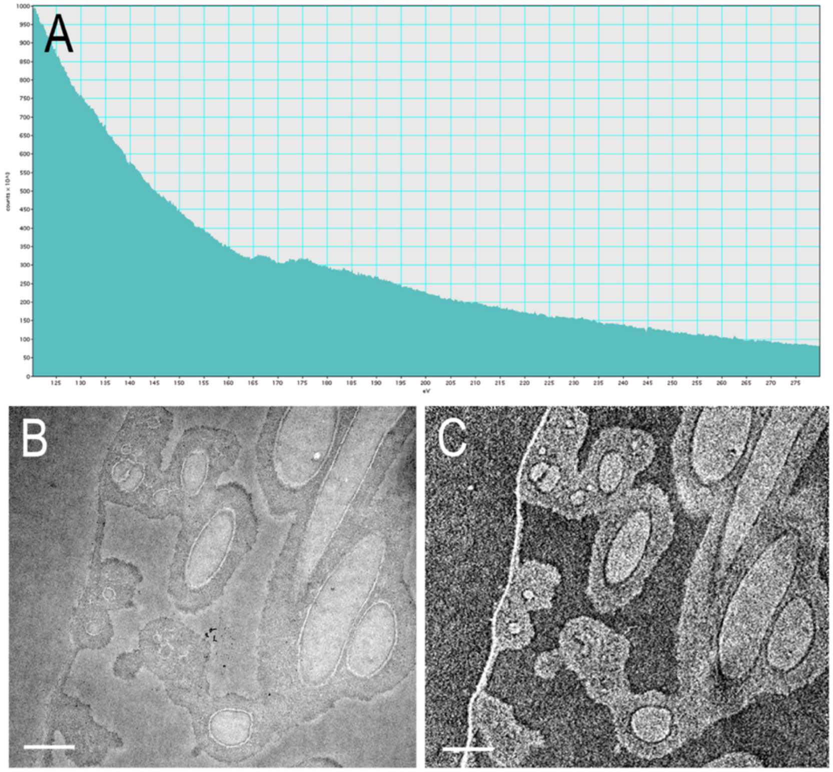

**Figure S12 | Electron Energy-Loss Spectroscopy (A), brightfield (B), and Energy-filtered TEM (C) images of eggshell membrane.** **Panel A**. A representative EELS spectrum acquired from the core region of a fiber, which shows the sulfur L2,3 edge at 161 eV. **Panel B**. Zero-loss brightfield image of eggshell membrane. **Panel C**. Elemental map of sulfur acquired with the three-window method. Bright areas indicate the presence of sulfur. Scale bars are one micron.

**Table SI | Summary of solution NMR experiments*.**

| Experiment | NS | NUS% | Number of points | | | Spectral width (ppm) | | |
|---|---|---|---|---|---|---|---|---|
| | | | F3 ($^{1}H$) | F2 ($^{15}N$) | F1 ($^{13}C$) | $^{1}H$ | $^{15}N$ | $^{13}C$ |
| | | | U-$^{13}C$,$^{15}N$ (10% $D_2O$) | | | | | |
| $^{15}N$-HSQC | 8 | 50 | 2048 | 512 | - | 19.8 | 50 | - |
| CBCA(CO)NH | 16 | 40 | 2048 | 86 | 256 | 16 | 36 | 60 |
| HNCO | 8 | 25 | 2048 | 120 | 180 | 16 | 36 | 12 |
| HNCA | 8 | 25 | 2048 | 84 | 160 | 13.65 | 36 | 31 |
| HNCACB | 24 | 25 | 2048 | 90 | 228 | 16 | 36 | 60 |
| CCCONH | 16 | 25 | 2048 | 96 | 196 | 16 | 36 | 60 |
| HNCACO | 32 | 25 | 2048 | 80 | 180 | 16 | 36 | 12 |
| HBCBCGCDCEHE | 256 | 50 | 2048 | - | 164 | 16 | - | 40 |
| HBCBCGCDHD | 48 | 50 | 2048 | - | 64 | 16 | - | 40 |
| | | | Number of points | | | Spectral width (ppm) | | |
| | | | F3 ($^{1}H$) | F2 ($^{15}N$) | F1 ($^{15}N$) | $^{1}H$ | $^{1}H$ | $^{15}N$ |
| HNHA | 16 | 25 | 2048 | 164 | 96 | 16 | 12 | 36 |
| HNHB | 32 | 25 | 2048 | 164 | 80 | 16 | 12 | 36 |
| HCCONH | 16 | 25 | 2048 | 80 | 164 | 16 | 36 | 16 |
| HBHACONH | 16 | 25 | 2048 | 96 | 180 | 16 | 36 | 11 |
| | | | U-$^{13}C$,$^{15}N$ (5% $D_2O$) | | | | | |
| | | | Number of points | | | Spectral width (ppm) | | |
| | | | F3 ($^{1}H$) | F2 ($^{15}N$) | F1 ($^{1}H$) | $^{1}H$ | $^{15}N$ | $^{1}H$ |
| $^{15}N$-edited NOESY | 32 | 25 | 2048 | 144 | 256 | 13.65 | 35 | 13.65 |
| | | | U-$^{13}C$,$^{15}N$ (100% $D_2O$) | | | | | |
| | | | Number of points | | | Spectral width (ppm) | | |
| | | | F3 ($^{1}H$) | F2 ($^{13}C$) | F1 ($^{1}H$) | $^{1}H$ | $^{13}C$ | $^{1}H$ |
| $^{13}C$-edited NOESY (Aliphatic) | 48 | 25 | 2048 | 64 | 160 | 15 | 24 | 12 |
| $^{13}C$-edited NOESY (Aromatic) | 48 | 25 | 2048 | 64 | 160 | 15 | 23 | 12 |

*All experiments were performed at 14.1 T and 298 K. The protein concentration was approximately 0.4 mM.

**Table SII | NMR chemical shifts for 4-mod CREMP**.

| Residue # | H | N | ND | NE | C | CA | CB | CG | CD | CE | CZ | CH |
|---|---|---|---|---|---|---|---|---|---|---|---|---|
| S1 | - | - | - | - | - | - | - | - | - | - | - | - |
| G2 | - | - | - | - | 174.22 | 45.24 | - | - | - | - | - | - |
| G3 | 8.43 | 108.99 | - | - | 175.06 | 45.57 | - | - | - | - | - | - |
| C4 | 8.64 | 118.74 | - | - | 175.6 | 57.26 | 39.88 | - | - | - | - | - |
| N5 | 8.35 | 117.11 | - | - | 175.47 | 55.14 | 38.04 | - | - | - | - | - |
| D6 | 7.84 | 116.75 | - | - | 175.15 | 54.91 | 41.88 | - | - | - | - | - |
| I7 | 7.02 | 115.95 | - | - | 174.2 | 59.18 | 40.36 | 26.85<br>17.54 | 14.25 | - | - | - |
| Y8 | 8.33 | 124.77 | - | - | 175.04 | 56.89 | 39.01 | - | 133.19 | 117.92 | - | - |
| C9 | 8.21 | 124.93 | - | - | 172.69 | 50.63 | 37.32 | - | - | - | - | - |
| P10 | - | - | - | - | 176.38 | 62.16 | 32.87 | 27.1 | 50.85 | - | - | - |
| K11 | 8.51 | 121.79 | - | - | 177.13 | 58.3 | 32.2 | 24.66 | 29.26 | 42.13 | - | - |
| G12 | 8.99 | 113.13 | - | - | 173.68 | 44.95 | - | - | - | - | - | - |
| T13 | 7.96 | 110.81 | - | - | 173.52 | 60.05 | 72.79 | 21.81 | - | - | - | - |
| T14 | 8.75 | 114.68 | - | - | 172.45 | 61.2 | 72.27 | 20.88 | - | - | - | - |
| C15 | 8.81 | 126.15 | - | - | 174.01 | 57.56 | 39.85 | - | - | - | - | - |
| K16 | 9.44 | 131.23 | - | - | 175.11 | 55.21 | 36.88 | 25.22 | 29.37 | 42.3 | - | - |
| M17 | 8.46 | 121.25 | - | - | 176.35 | 53.44 | 30.15 | 31.72 | - | 17.58 | - | - |
| M18 | 8.64 | 125.35 | - | - | 175.12 | 54.28 | 34.16 | 31.5 | - | - | - | - |
| D19 | 9.23 | 125.37 | - | - | 175.95 | 55.47 | 39.88 | - | - | - | - | - |
| G20 | 8.42 | 101.64 | - | - | 173.41 | 45.22 | - | - | - | - | - | - |
| H21 | 7.76 | 120.86 | - | - | 173.29 | 53.89 | 32.03 | - | 122.2 | 136.33 | - | - |
| P22 | - | - | - | - | 176.48 | 63.05 | 32.07 | 27.37 | 50.7 | - | - | - |
| Q23 | 8.78 | 122.83 | - | - | 174.04 | 54.55 | 32.89 | 33.68 | - | - | - | - |
| C24 | 8.87 | 121.94 | - | - | 174.79 | 53.53 | 37.48 | - | - | - | - | - |

| | | | | | | | | | | | | |
|---|---|---|---|---|---|---|---|---|---|---|---|---|
| V25 | 8.93 | 125.15 | - | - | 175.79 | 60.91 | 34.17 | 21.44<br>20.87 | - | - | - | - |
| Q26 | 8.78 | 125.7 | - | - | 175.78 | 57.21 | 29.2 | 33.68 | - | - | - | - |
| N27 | 8.3 | 123.46 | 109.85 | - | 175.22 | 53.4 | 38.35 | - | - | - | - | - |
| G28 | 8.33 | 121.83 | - | - | 173.35 | 56 | 28.94 | - | - | - | - | - |
| P29 | - | - | - | - | 175.49 | 63.07 | 32.7 | 27.25 | 50.65 | - | - | - |
| S30 | 8.61 | 115.87 | - | - | 175.18 | 57.43 | 66.52 | - | - | - | - | - |
| C31 | 8.76 | 118.66 | - | - | 175.73 | 57.85 | 39.62 | - | - | - | - | - |
| S32 | 8.13 | 114.47 | - | - | 174.7 | 60.61 | 62.85 | - | - | - | - | - |
| D33 | 7.71 | 118.04 | - | - | 175.55 | 55.44 | 42.63 | - | - | - | - | - |
| I34 | 7.1 | 114.17 | - | - | 173.82 | 59.25 | 41.39 | 17.69<br>26.54 | 14.32 | - | - | - |
| Q35 | 8.22 | 121.54 | - | - | 175.47 | 54.29 | 30.21 | 33.83 | - | - | - | - |
| C36 | 8.76 | 123.26 | - | - | 175.28 | 52.7 | 38.18 | - | - | - | - | - |
| S37 | 8.24 | 118.31 | - | - | 174.24 | 58.35 | 64.21 | - | - | - | - | - |
| K38 | 8.58 | 122.62 | - | - | 177.17 | 58.37 | 32.03 | 24.65 | 29.36 | 42.13 | - | - |
| G39 | 8.91 | 112.7 | - | - | 173.99 | 44.98 | - | - | - | - | - | - |
| T40 | 8.01 | 111.15 | - | - | 173.02 | 60.08 | 72.71 | 22.05 | - | - | - | - |
| T41 | 8.75 | 113.1 | - | - | 172.44 | 60.45 | 71.51 | 20.72 | - | - | - | - |
| C42 | 8.73 | 125.58 | - | - | 174.34 | 57.34 | 40.73 | - | - | - | - | - |
| Q43 | 9.34 | 129.94 | - | - | 174.45 | 54.73 | 33.01 | 33.85 | - | - | - | - |
| M44 | 8.58 | 121.62 | - | - | 176.67 | 53.39 | 29.86 | 31.6 | - | - | - | - |
| V45 | 9.05 | 127.25 | - | - | 175.88 | 61.98 | 34.01 | 20.68 | - | - | - | - |
| D46 | 9.37 | 128.52 | - | - | 175.85 | 55.49 | 39.85 | - | - | - | - | - |
| G47 | 8.42 | 101.62 | - | - | 173.8 | 45.28 | - | - | - | - | - | - |
| W48 | 7.74 | 117.97 | - | 128.84 | 171.61 | 52.55 | 30.3 | 109.23 | 127.96 | 120.5 | 114.62<br>121.14 | 124.61 |
| P49 | - | - | - | - | 175.98 | 62.85 | 31.67 | 27.21 | 50.71 | - | - | - |
| R50 | 8.88 | 122.76 | - | - | 174.49 | 53.93 | 34.26 | 26.77 | 43.35 | - | - | - |
| C51 | 8.7 | 120.52 | - | - | 174.76 | 53.76 | 38.43 | - | - | - | - | - |

| | | | | | | | | | | | | |
|---|---|---|---|---|---|---|---|---|---|---|---|---|
| I52 | 9.1 | 125.69 | - | - | 175.78 | 58.75 | 40 | 26.53<br>17.57 | 12.05 | - | - | - |
| Q53 | 8.65 | 125.18 | - | - | 175.86 | 56.88 | 29.17 | 33.9 | - | - | - | - |
| I54 | 7.99 | 124.85 | - | - | 176.09 | 61.96 | 38.85 | 27.91<br>17.27 | 13.4 | - | - | - |
| K55 | 8.33 | 126.08 | - | - | 175.68 | 56 | 33.19 | 24.78 | 29.2 | 42.08 | - | - |
| A56 | 8.22 | 126.28 | - | - | 176.92 | 52.11 | 19.49 | - | - | - | - | - |
| A57 | 8.23 | 125.13 | - | - | 175.46 | 50.38 | 18.18 | - | - | - | - | - |
| P58 | - | - | - | - | 176.89 | 62.9 | 32.11 | 27.41 | 50.63 | - | - | - |
| S59 | 8.34 | 116.68 | - | - | 174.22 | 58.37 | 63.85 | - | - | - | - | - |
| R60 | 8.3 | 123.49 | - | - | 173.85 | 53.82 | 30.24 | - | - | - | - | - |
| P61 | - | - | - | - | 175.84 | 63.24 | 32.71 | 27.57 | 50.66 | - | - | - |
| S62 | 8.77 | 116.75 | - | - | 175.18 | 57.42 | 66.41 | - | - | - | - | - |
| C63 | 8.68 | 118.53 | - | - | 175.82 | 57.95 | 39.57 | - | - | - | - | - |
| S64 | 8.17 | 114.61 | - | - | 174.83 | 60.61 | 62.97 | - | - | - | - | - |
| D65 | 7.73 | 118.35 | - | - | 175.48 | 55.43 | 42.63 | - | - | - | - | - |
| I66 | 7.03 | 114.44 | - | - | 173.87 | 59.23 | 41.16 | 17.73 | - | - | - | - |
| H67 | - | - | - | - | 174.64 | 53.93 | 29.54 | - | 119.85 | 136.61 | - | - |
| C68 | 8.86 | 124.75 | - | - | 173.36 | 51.88 | 36.88 | - | - | - | - | - |
| P69 | - | - | - | - | 176.24 | 61.88 | 32.91 | 27.11 | 50.77 | - | - | - |
| K70 | 8.42 | 121.4 | - | - | 177.13 | 58.28 | 32.18 | 24.72 | 29.28 | 42.11 | - | - |
| G71 | 8.99 | 113.13 | - | - | 173.67 | 44.96 | - | - | - | - | - | - |
| T72 | 7.99 | 111.13 | - | - | 173.53 | 60.06 | 72.79 | 22.06 | - | - | - | - |
| T73 | 8.78 | 114.68 | - | - | 172.45 | 61.2 | 72.27 | 20.88 | - | - | - | - |
| C74 | 8.81 | 126.13 | - | - | 174.01 | 57.56 | 40.61 | - | - | - | - | - |
| K75 | 9.45 | 131.3 | - | - | 174.86 | 55.12 | 36.9 | 25.13 | 29.25 | 42.29 | - | - |
| M76 | 8.43 | 120.83 | - | - | 176.02 | 53.13 | 29.78 | 31.67 | - | - | - | - |
| I77 | 8.63 | 127.54 | - | - | 175.83 | 59.89 | 39.53 | 26.55<br>16.87 | 12.42 | - | - | - |
| D78 | 9.3 | 128.27 | - | - | 176.08 | 55.5 | 39.78 | - | - | - | - | - |

| | | | | | | | | | | | | |
|---|---|---|---|---|---|---|---|---|---|---|---|---|
| G79 | 8.43 | 101.56 | - | - | 173.34 | 45.25 | - | - | - | - | - | - |
| H80 | 7.8 | 121.34 | - | - | 173.47 | 53.87 | 32.06 | - | 122.18 | 136.33 | - | - |
| P81 | - | - | - | - | 176.4 | 62.9 | 31.76 | 27.27 | 50.71 | - | - | - |
| Q82 | 8.95 | 123.12 | - | - | 174.06 | 54.52 | 32.74 | 33.78 | - | - | - | - |
| C83 | 8.87 | 121.88 | - | - | 174.77 | 53.53 | 37.59 | - | - | - | - | - |
| V84 | 8.94 | 125.15 | - | - | 175.79 | 60.9 | 34.17 | 20.82<br>21.35 | - | - | - | - |
| Q85 | 8.79 | 125.73 | - | - | 175.77 | 57.23 | 29.22 | 33.73 | - | - | - | - |
| N86 | 8.3 | 123.47 | - | - | 175.17 | 53.4 | 38.34 | - | - | - | - | - |
| Q87 | 8.32 | 121.86 | - | - | 173.28 | 56 | 28.92 | - | - | - | - | - |
| P88 | - | - | - | - | 175.76 | 62.96 | 32.6 | 27.13 | 50.68 | - | - | - |
| T89 | 8.67 | 110.43 | - | - | 175.97 | 60.1 | 72.31 | 21.02 | - | - | - | - |
| C90 | 8.5 | 117.39 | - | - | 175.99 | 57.52 | 39.04 | - | - | - | - | - |
| S91 | 8.05 | 114.29 | - | - | 174.7 | 60.61 | 62.87 | - | - | - | - | - |
| D92 | 7.7 | 118.03 | - | - | 175.46 | 55.38 | 42.96 | - | - | - | - | - |
| I93 | 7.08 | 114.81 | - | - | 173.96 | 59.27 | 41.26 | 26.58<br>17.68 | 14.25 | - | - | - |
| Q94 | 8.23 | 121.59 | - | - | 175.47 | 54.29 | 30.2 | 33.91 | - | - | - | - |
| C95 | 8.76 | 123.28 | - | - | 175.33 | 52.7 | 38.08 | - | - | - | - | - |
| S96 | 8.2 | 118.37 | - | - | 174.26 | 58.31 | 64.2 | - | - | - | - | - |
| K97 | 8.6 | 122.64 | - | - | 177.18 | 58.4 | 32.08 | 24.6 | 29.36 | 42.19 | - | - |
| G98 | 8.95 | 112.84 | - | - | 173.87 | 44.95 | - | - | - | - | - | - |
| T99 | 8.06 | 111.15 | - | - | 172.84 | 59.98 | 72.89 | 21.93 | - | - | - | - |
| T100 | 8.79 | 113.28 | - | - | 172.22 | 60.39 | 71.39 | 20.48 | - | - | - | - |
| C101 | 8.63 | 125.43 | - | - | 174.59 | 57.28 | 41.28 | - | - | - | - | - |
| Q102 | 9.5 | 130.8 | - | - | 174.36 | 54.72 | 32.99 | 33.87 | - | - | - | - |
| M103 | 8.6 | 121.48 | - | - | 176.56 | 53.34 | 29.94 | 31.67 | - | - | - | - |
| V104 | 8.99 | 127.21 | - | - | 175.87 | 61.97 | 34.02 | 20.79 | - | - | - | - |
| D105 | 9.37 | 128.53 | - | - | 175.76 | 55.52 | 39.87 | - | - | - | - | - |

| | | | | | | | | | | | | |
|---|---|---|---|---|---|---|---|---|---|---|---|---|
| G106 | 8.42 | 101.61 | - | - | 173.8 | 45.28 | - | - | - | - | - | - |
| W107 | 7.79 | 117.93 | - | 128.93 | 171.54 | 52.64 | 30.79 | - | 127.88 | 120.59 | 114.66<br>121.18 | 124.62 |
| P108 | - | - | - | - | 176.02 | 62.87 | 31.62 | 27.23 | 50.71 | - | - | - |
| R109 | 8.97 | 122.64 | - | - | 175 | 54.29 | 33.88 | 26.51 | 43.53 | - | - | - |
| C110 | 8.83 | 122.08 | - | - | 175.35 | 54.08 | 38.6 | - | - | - | - | - |
| T111 | 9.13 | 119.71 | - | - | 173.42 | 60.78 | 71.12 | 21.15 | - | - | - | - |
| Q112 | 8.59 | 122.79 | - | - | 175.91 | 56.43 | 29.56 | 33.93 | - | - | - | - |
| L113 | 8.13 | 125.53 | - | - | 176.94 | 55.37 | 42.62 | 27.19 | 24.85<br>23.85 | - | - | - |
| K114 | 8.37 | 123.52 | - | - | 176.13 | 56.11 | 33.09 | 24.86 | 29.19 | 42.08 | - | - |
| V115 | 8.05 | 122.21 | - | - | 175.36 | 61.69 | 33.08 | 21.20<br>20.49 | - | - | - | - |
| A116 | 8.36 | 129.97 | - | - | 176.23 | 50.35 | 18.17 | - | - | - | - | - |
| P117 | - | - | - | - | 176.7 | 62.81 | 32.21 | 27.44 | 50.58 | - | - | - |
| R118 | 8.32 | 121.77 | - | - | 176.15 | 56 | 30.96 | 26.98 | 43.33 | - | - | - |
| R119 | 8.38 | 124.25 | - | - | 174.33 | 53.93 | 30.35 | - | 43.33 | - | - | - |
| P120 | - | - | - | - | 176.11 | 63.58 | 32.09 | 27.36 | 50.78 | - | - | - |
| S121 | 7.88 | 121.63 | - | - | 178.6 | 60.04 | 64.84 | - | - | - | - | - |

**Table SII (con'd) | NMR chemical shifts for 4-mod CREMP**.

| Residue # | HA | HB | HG | HD | HE | HZ | HH |
|---|---|---|---|---|---|---|---|
| S1 | 4.79 | - | 4.71 | - | - | - | - |
| G2 | 3.99<br>3.45 | - | - | - | - | - | - |
| G3 | 4.15<br>3.88 | - | - | - | - | - | - |
| C4 | 4.45 | 2.79<br>3.02 | - | - | - | - | - |
| N5 | 4.46 | 2.48<br>2.70 | - | - | - | - | - |
| D6 | 4.54 | 2.79<br>2.61 | - | - | - | - | - |
| I7 | 4.41 | 1.82 | 0.69<br>1.23<br>1.04 | 0.62 | - | - | - |
| Y8 | 4.61 | 2.80<br>2.85 | - | 7.09 | 6.8 | - | - |
| C9 | 5.11 | 3.59<br>2.12 | - | - | - | - | - |
| P10 | 4.44 | 2.27<br>1.92 | 2.02<br>1.85 | 3.64 | - | - | - |
| K11 | 4.07 | 1.78<br>1.75 | 1.44<br>1.34 | 1.57<br>1.68 | 2.99 | - | - |
| G12 | 4.30<br>3.62 | - | - | - | - | - | - |
| T13 | 4.97 | 3.5 | 0.78 | - | - | - | - |
| T14 | 4.39 | 3.86 | 1.11 | - | - | - | - |
| C15 | 4.96 | 3.07<br>2.84 | - | - | - | - | - |
| K16 | 4.5 | 1.67<br>1.57 | 1.29<br>1.36 | 1.53<br>1.60 | 2.82<br>2.91 | - | - |
| M17 | 4.69 | 1.88<br>1.69 | 2.33<br>2.55 | - | 0.83 | - | - |
| M18 | 4.62 | 1.91<br>1.76 | 2.52<br>2.34 | - | 0.82 | - | - |
| D19 | 4.16 | 2.91<br>2.88 | - | - | - | - | - |
| G20 | 3.98<br>3.35 | - | - | - | - | - | - |
| H21 | 4.88 | 2.86<br>3.20 | - | 7.18 | 8.61<br>9.06 | - | - |
| P22 | 4.62 | 1.68<br>1.80 | 2.02<br>2.00 | 3.8 | - | - | - |
| Q23 | 4.52 | 1.97<br>1.79 | 2.49<br>2.24 | - | - | - | - |
| C24 | 4.85 | 2.73<br>2.90 | - | - | - | - | - |
| V25 | 4.33 | 2.01 | 0.87<br>0.63 | - | - | - | - |

| | | | | | | | |
|---|---|---|---|---|---|---|---|
| Q26 | 4.02 | 2.00<br>1.83 | 2.36<br>2.42 | - | - | - | - |
| N27 | 4.5 | 2.62<br>2.53 | - | 6.67<br>7.45 | - | - | - |
| G28 | 4.41 | 1.93<br>1.74 | 2.19 | - | 7.45<br>6.66 | - | - |
| P29 | 4.43 | 2.08<br>1.54 | 1.51<br>1.66 | 3.56<br>3.37 | - | - | - |
| S30 | 4.79 | 3.70<br>3.98 | - | - | - | - | - |
| C31 | 4.46 | 3.01<br>2.85 | - | - | - | - | - |
| S32 | 4.28 | 3.75<br>3.67 | 4.68 | - | - | - | - |
| D33 | 4.55 | 2.71<br>2.82 | - | - | - | - | - |
| I34 | 4.52 | 1.8 | 1.22<br>0.76<br>1.00 | 0.64 | - | - | - |
| Q35 | 4.55 | 1.89<br>1.93 | 2.24<br>2.30 | - | - | - | - |
| C36 | 4.92 | 3.64<br>2.25 | - | - | - | - | - |
| S37 | 4.32 | 3.83<br>3.64 | 4.72 | - | - | - | - |
| K38 | 4.07 | 1.75<br>1.64 | 1.43 | 1.69 | 2.99 | - | - |
| G39 | 3.63<br>4.29 | - | - | - | - | - | - |
| T40 | 4.89 | 3.78 | 0.79 | - | - | - | - |
| T41 | 4.5 | 3.87 | 1.04 | - | - | - | - |
| C42 | 5 | 3.15<br>2.84 | - | - | - | - | - |
| Q43 | 4.54 | 1.73<br>1.95 | 2.25<br>2.29 | - | 6.73<br>7.17 | - | - |
| M44 | 4.76 | 1.68<br>1.93 | 2.37<br>2.55 | - | 1.95 | - | - |
| V45 | 4.14 | 1.93 | 0.83<br>0.88 | - | - | - | - |
| D46 | 4.18 | 2.61<br>2.96 | - | - | - | - | - |
| G47 | 3.98<br>3.36 | - | - | - | - | - | - |
| W48 | 5.06 | 3.01<br>3.17 | - | 7.25 | 9.99<br>7.77 | 7.30<br>7.51 | 7.27 |
| P49 | 4.62 | 1.70<br>1.56 | 1.91 | 3.77<br>3.51 | - | - | - |
| R50 | 4.58 | 1.73 | 1.66 | 3.19<br>3.02 | - | - | - |
| C51 | 4.88 | 2.90<br>2.69 | - | - | - | - | - |
| I52 | 4.44 | 1.89 | 1.30<br>1.26<br>0.86 | 0.68 | - | - | - |

| | | | | | | | |
|---|---|---|---|---|---|---|---|
| Q53 | 4.19 | 1.95 | 2.37<br>2.31 | - | - | - | - |
| I54 | 3.94 | 1.64 | 1.34<br>0.82<br>1.29 | 0.76 | - | - | - |
| K55 | 4.25 | 1.71<br>1.65 | 1.41<br>1.35 | 1.58<br>1.67 | 2.97 | - | - |
| A56 | 4.23 | 1.31 | - | - | - | - | - |
| A57 | 4.56 | 1.3 | - | - | - | - | - |
| P58 | 4.39 | 2.26<br>1.84 | 1.99<br>1.86 | 3.63<br>3.80 | - | - | - |
| S59 | 4.31 | 3.77 | - | - | - | - | - |
| R60 | 4.6 | 1.79<br>1.95 | 1.62 | 3.23 | 7.45 | - | 6.77<br>6.67 |
| P61 | 4.5 | 2.23<br>1.72 | 1.95 | 3.74 | - | - | - |
| S62 | 4.8 | 3.77<br>4.03 | - | - | - | - | - |
| C63 | 4.43 | 2.99<br>2.84 | - | - | - | - | - |
| S64 | 4.27 | 3.82<br>3.74 | 4.75 | - | - | - | - |
| D65 | 4.5 | 2.73<br>2.85 | - | - | - | - | - |
| I66 | 4.44 | 1.76 | 0.73<br>1.21<br>1.53 | 0.64 | - | - | - |
| H67 | 4.74 | 2.99<br>2.97 | - | 7.19 | 8.46 | - | - |
| C68 | 4.78 | 3 | - | - | - | - | - |
| P69 | 4.31 | 2.23<br>1.97 | 1.88<br>1.83 | 3.80<br>3.65 | - | - | - |
| K70 | 4.05 | 1.76<br>1.75 | 1.36<br>1.43 | 1.68 | 2.99 | - | - |
| G71 | 4.29<br>3.60 | - | - | - | - | - | - |
| T72 | 4.96 | 3.5 | 0.78 | - | - | - | - |
| T73 | 4.39 | 3.85 | 1.11 | - | - | - | - |
| C74 | 5.01 | 2.83<br>3.10 | - | - | - | - | - |
| K75 | 4.53 | 1.70<br>1.56 | 1.34 | 1.62 | 2.87 | - | - |
| M76 | 4.71 | 1.64<br>1.93 | 2.34<br>2.58 | - | - | - | - |
| I77 | 4.22 | 1.67 | 0.76<br>1.02<br>1.25 | 0.64 | - | - | - |
| D78 | 4.21 | 2.98<br>2.59 | - | - | - | - | - |
| G79 | 4.02<br>3.36 | - | - | - | - | - | - |
| H80 | 4.92 | 3.21 | - | 7.19 | 8.59 | - | - |

| | | | | | | | |
|---|---|---|---|---|---|---|---|
| | | 2.92 | | | 9.0 | | |
| P81 | 4.64 | 1.74<br>1.82 | 1.98<br>2.07 | 3.54<br>3.81 | - | - | - |
| Q82 | 4.56 | 1.98<br>1.81 | 2.29<br>2.24 | - | - | - | - |
| C83 | 4.84 | 2.90<br>2.73 | - | - | - | - | - |
| V84 | 4.37 | 2.05 | 0.89<br>0.97 | - | - | - | - |
| Q85 | 3.99 | 1.98<br>1.78 | 2.48<br>2.37 | - | - | - | - |
| N86 | 4.32 | 2.53<br>2.66 | - | 7.46<br>6.67 | - | - | - |
| Q87 | 4.41 | 1.91<br>1.74 | 2.37 | - | 7.45<br>6.67 | - | - |
| P88 | 4.48 | 2.12<br>1.58 | 1.52<br>1.67 | 3.62<br>3.54 | - | - | - |
| T89 | 4.68 | 4.42 | 1.09 | - | - | - | - |
| C90 | 4.48 | 2.85<br>2.98 | - | - | - | - | - |
| S91 | 4.3 | 3.75<br>3.64 | - | - | - | - | - |
| D92 | 4.55 | 2.72<br>2.82 | - | - | - | - | - |
| I93 | 4.5 | 1.79 | 1.04<br>1.22<br>0.73 | 0.64 | - | - | - |
| Q94 | 4.54 | 1.88 | 2.22 | - | - | - | - |
| C95 | 4.92 | 3.69<br>2.26 | - | - | - | - | - |
| S96 | 4.33 | 3.83<br>3.65 | 4.72 | - | - | - | - |
| K97 | 4.06 | 1.72<br>1.75 | 1.43 | 1.58 | - | - | - |
| G98 | 4.33<br>3.65 | - | - | - | - | - | - |
| T99 | 5.08 | 3.61 | 0.82 | - | - | - | - |
| T100 | 4.48 | 3.86 | 1.05 | - | - | - | - |
| C101 | 5.09 | 2.83<br>3.16 | - | - | - | - | - |
| Q102 | 4.54 | 1.93<br>1.78 | 2.29<br>2.21 | - | 7.17 | - | - |
| M103 | 4.77 | 1.67<br>1.92 | 2.55<br>2.34 | - | - | - | - |
| V104 | 4.14 | 1.91 | 0.80<br>0.89 | - | - | - | - |
| D105 | 4.21 | 2.68<br>2.99 | - | - | - | - | - |
| G106 | 3.37<br>3.99 | - | - | - | - | - | - |
| W107 | 5 | 3.12<br>3.04 | - | 7.27 | 7.77<br>10.00 | 7.29<br>7.52 | 7.27 |

| | | | | | | | |
|---|---|---|---|---|---|---|---|
| P108 | 4.68 | 1.84<br>1.74 | 2.06<br>2.00 | 3.53<br>3.80 | - | - | - |
| R109 | 4.57 | 1.77<br>1.81 | 1.44<br>1.67 | 3.18<br>3.04 | - | - | - |
| C110 | 4.95 | 2.92<br>2.71 | - | - | - | - | - |
| T111 | 4.52 | 4.07 | 1.08 | - | - | - | - |
| Q112 | 4.26 | 2.03<br>1.95 | 2.34 | - | - | - | - |
| L113 | 4.2 | 1.46 | 1.44 | 0.81<br>0.83 | - | - | - |
| K114 | 4.29 | 1.71 | 1.40<br>1.32 | - | 2.96 | - | - |
| V115 | 4.04 | 1.96 | 0.86<br>0.90 | - | - | - | - |
| A116 | 4.56 | 1.3 | - | - | - | - | - |
| P117 | 4.33 | 1.80<br>2.22 | 1.95 | - | - | - | - |
| R118 | 4.24 | 1.87<br>1.74 | 1.63<br>1.53 | 3.18<br>3.26 | - | - | - |
| R119 | 4.59 | 1.79<br>1.69 | - | 3.19 | - | - | - |
| P120 | 4.41 | 1.97<br>2.28 | 2.03 | 3.63<br>3.80 | - | - | - |
| S121 | 4.16 | 3.93<br>3.79 | - | - | - | - | - |

**Table SIII | Intra-module distance restraints.**

| | Mod1 (1-30) | Mod2 (31-62) | Mod3 (63-89) | Mod4 (90-121) | Total |
|---|---|---|---|---|---|
| **Intra-residue** | 87 (2.9)* | 100 (3.1) | 78 (3) | 93 (3) | 358 (2.9) |
| **Sequential** | 129 (4.3) | 155 (4.8) | 118 (4.5) | 132 (4.2) | 534 (4.4) |
| **Medium-range** | 78 (2.6) | 90 (2.8) | 63 (2.4) | 76 (2.4) | 307 (2.5) |
| **Long-range** | 72 (2.4) | 27 (0.8) | 61 (2.3) | 43 (1.3) | 203 (1.6) |
| **Total** | 366 | 372 | 320 | 344 | 1402 |

* Total restraints (restraints per residue)

**Table SIV | Inter-module distance restraints.**

| | Mod1-2 | Mod2-3 | Mod3-4 | Total |
|---|---|---|---|---|
| **Intra-residue** | - | - | - | - |
| **Sequential** | 7 | 5 | 5 | 17 |
| **Medium-range (resd if 2-4)** | 15 | 5 | 8 | 28 |
| **Long-range (resd if >4)** | - | - | 1 | 1 |
| **Total** | 22 | 10 | 14 | 46 |
